# Developmental evolution of social signals in the delayed plumage maturation of manakins (Aves: Pipridae)

**DOI:** 10.1101/2025.05.15.654367

**Authors:** Liam U. Taylor, Richard Owen Prum

## Abstract

Avian plumage maturation involves replacing feathers, via discrete molts, until reaching an iteratively-regenerated definitive plumage. In most birds, this process takes about one year. In the Neotropical lekking manakins (Pipridae), males of most species exhibit delayed plumage maturation (DPM), passing through drab predefinitive plumages for two or three years before reaching a bright, sexually dimorphic, definitive plumage. We used a phylogenetic analysis to investigate the evolutionary history of DPM in manakins. Our unique comparative dataset derived characters from a graph structure representing the developmental schedules of individual, colorful plumage patches. We traced the origin of one-year DPM to the ancestor of sexually dichromatic Piprinae manakins, followed by three separate origins of prolonged, two-year DPM, plus one subsequent extension to three-year DPM. Consistent with social signaling hypotheses, prolonged DPM evolved to reveal and maintain signals of young male status. This interspecific pattern sheds light on the sociosexual development of lekking birds, a topic usually restricted to autecological field studies. By detailing heterochronic dynamics in manakin plumage evolution, we also show how the iterative regeneration of characters like bird feathers, insect instars, mammalian antlers, and angiosperm flowers can create multiple distinct levels of developmental homologies among bodies, sexes, and species.

## Introduction

Comparative analyses of developmental evolution traditionally assumed that development is a continuous process of growth or accumulation (Gould 1977). In this view, developmental evolution can be neatly described in terms of start times, stop times, or growth rates for simple, universally homologous characters (e.g., body size, univariate body shape; Alberch et al. 1979; Fink 1982). This approach fails to describe the true diversity of development. In reality, developmental evolution can include shifts in the form of discrete characters (Wake and Larson 1987), changes in the ontogenetic order or schedule of those characters (Bininda-Emonds et al. 2002; Smith 2002; Jeffery et al. 2005), the origination of entirely new characters (McKenna et al. 2021), and the iterative regeneration of characters over the life cycle (this study).

These developmental complexities are all apparent in avian plumage maturation, the process through which birds obtain their “definitive” adult plumage. Birds replace old feathers with new ones through a series of periodic molts (Dwight 1900; Payne 1972). A newly developed plumage can look identical to, or completely different from, the preceding plumage. Across successive molts, young birds undergo a broader progression in feather form and coloration. Plumage maturation usually begins with a downy natal plumage replaced within weeks or months by a loosely-textured juvenile plumage (Butler et al. 2008). Many birds then achieve their mature, or “definitive,” plumage through one or more molts in their first year of life (Humphrey and Parkes 1959; Howell et al. 2003; Wolfe et al. 2014). This definitive plumage is then regenerated via annual or semi-annual molts for the remaining life of the bird. Thus, avian plumage maturation is both a discrete and iterative process, the evolution of which can involve changes to both the form of ontogenetic characters (e.g., the characteristics of a plumage) and the schedule of those characters during development (e.g., the timing of acquisition or retention of those plumages).

Manakins (Passeriformes: Pipridae) are Neotropical lekking birds that exhibit especially complicated forms of delayed plumage maturation (DPM; Kirwan and Green 2011). Across the ∼60 manakin species, female definitive plumages are generally green with diffuse accents of olive, gray, or yellow. Male definitive plumages in most, though not all, species feature a diverse array of bright, colorful patches (Prum 1997; Doucet et al. 2007b; Ribeiro et al. 2015). Whereas female manakins reach definitive plumage at approximately one year old, males in different sexually dichromatic species take between two and four years to molt through a series of “predefinitive” plumages (Schaedler et al. 2021b). Such prolonged DPM is only known in a few, ecologically diverse lineages of birds, including some gulls (Charadriiformes: Laridae), albatrosses (Procellariiformes: Diomedeidae), raptors (Accipitriformes: Accipitridae), male bowerbirds (Passeriformes: Ptilonorhynchidae), and male birds of paradise (Passeriformes: Paradisaeidae; Hawkins et al. 2012).

Schaedler et al. (2021b) review the literature on manakin DPM, arguing that interspecific patterns of plumage evolution can shed light on the ecological function of predefinitive plumages. They note that multiple species of manakins have predefinitive plumage stages featuring potential signals of male status (Fig. 1). For example, in White-throated Manakins (*Corapipo gutturalis*), the male definitive plumage is black with a white throat. Young males in this species spend their first year in a green, female plumage, but then molt into a second predefinitive plumage with a black mask and white throat (Fig. 1B; Prum 1986).

**Figure 1.**
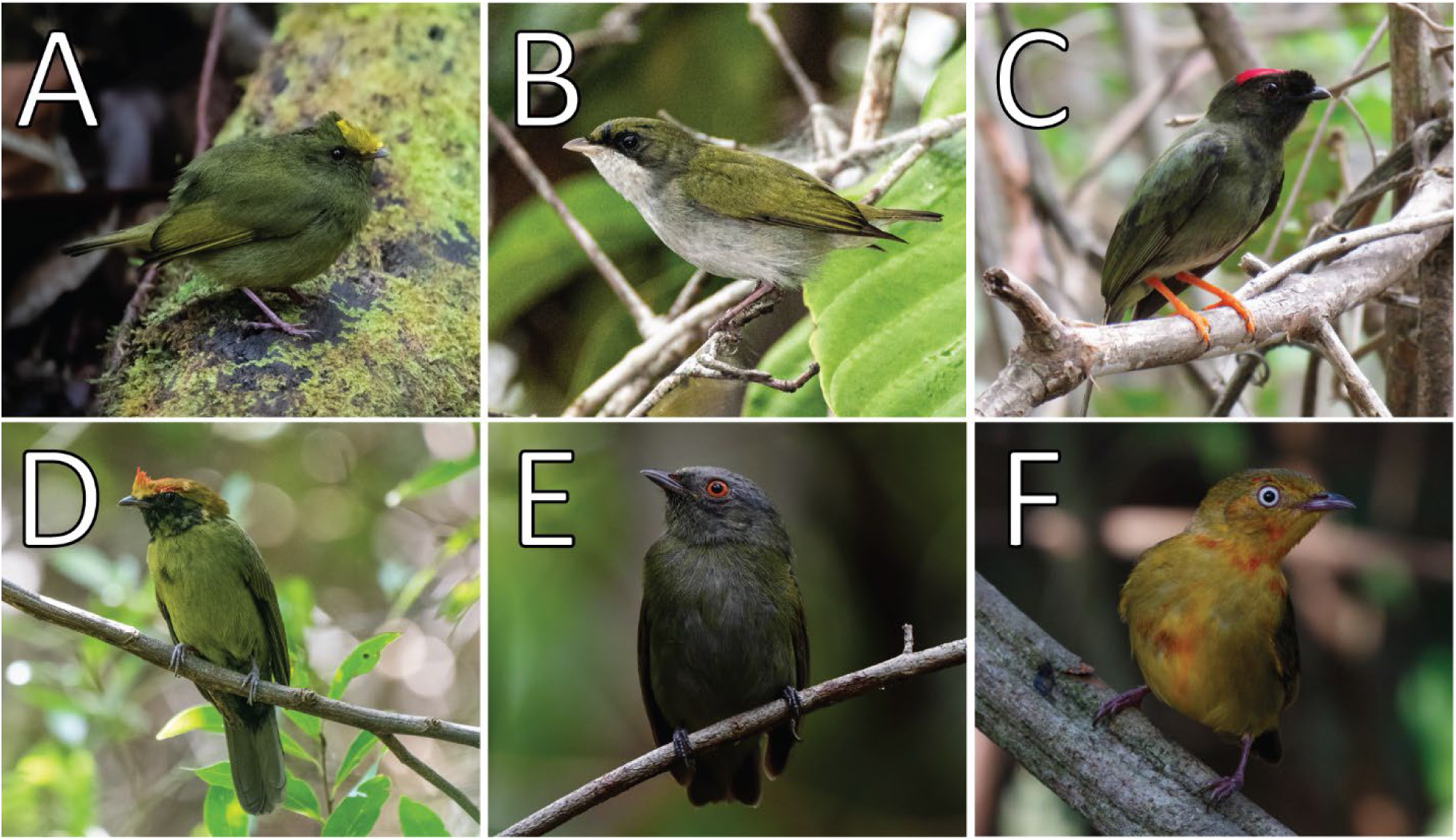
Examples of predefinitive plumages that feature distinct signals of young male status in six genera of manakins. All photographs reproduced with permission from Macaulay Library (ML). (A) *Masius chrysopterus* (ML608667835). (B) *Corapipo gutturalis* (ML363146471). (C) *Chiroxiphia linearis* (ML156641571). (D) *Antilophia galeata* (ML421373261). (E) *Pseudopipra pipra* (ML602227891). (F) *Pipra fasciicauda* (ML554623261).

The evolution of these male predefinitive plumage patches may be consistent with a social signaling hypothesis for DPM (Lyon and Montgomerie 1986; Hawkins et al. 2012). For lekking species like manakins, the social signaling hypothesis argues that predefinitive plumages evolve to help young birds indicate their status as inexperienced, less competitive males (Collis and Borgia 1993; McDonald 1993a; Prum and Razafindratsita 1997). These signals may then reduce the threat of competition from older individuals while young males integrate into the complex social environment of the lek and develop sociosexual relationships with other males. The social signaling hypothesis contrasts with alternative hypotheses regarding crypsis (Rohwer 1978) and female mimicry (Rohwer et al. 1980), which argue that predefinitive plumages evolve to obscure young male status, rather than reveal it (Hawkins et al. 2012; Schaedler et al. 2021b).

Here, we test the social signaling hypothesis with a phylogenetic, comparative analysis of manakin DPM. We ask: (1) whether DPM evolved to reveal, or obscure, signals of young male status relative to ancestral conditions; and (2) how predefinitive plumages maintain their social signaling properties when male identity itself diversifies across lineages of manakins. The social signaling hypothesis predicts: (1) DPM evolves to reveal young male status; and (2) the properties of predefinitive plumages must continue to evolve to *maintain* their function as salient, social signals of maleness.

To test these predictions, we modeled the phylogenetic history of manakin plumage maturation. Elaborating on previous phylogenetic analyses of avian DPM (Chu 1994; Hill 1996), we created a unique comparative dataset that incorporates both the properties of predefinitive plumages (i.e., colorful patches) and the developmental schedule of those plumages (i.e., when does a patch first appear, and how many times is it regenerated?). We found multiple origins of prolonged DPM in manakins, each evolving to reveal young male status, and each continuing to evolve to maintain signals of maleness. We discuss these results in terms of both behavioral ecology and developmental evolution.

## Materials and Methods

### Plumage maturation characters

Plumage maturation in manakins involves an iterative sequence of individual plumages, developed via discrete molts, that eventually converges on a definitive plumage. A phylogenetic analysis of plumage maturation requires a definition of characters and character states that leaves room for evolutionary changes in both individual plumage patches and the developmental schedules of those patches (see details in *Supplementary Material: Deriving characters from graph structures*). Species may differ in the aspect of male plumage at any point along the developmental schedule (e.g., *Pipra filicauda* and *P. fasciicauda* exhibit different plumages on a similar schedule; Ryder and Durães 2005; Scholer et al. 2021). Species may also differ in developmental schedule (e.g., *Chiroxiphia lanceolata* and *C. linearis* exhibit similar plumages on a different schedule; DuVal 2005; Doucet et al. 2007a). Here, we present a dataset that tracks evolutionary change to both plumage and developmental schedule across species.

We coded male plumage maturation in 40 manakin taxa for which sufficient data were available (Fig. 2). Full details, sources, and Macaulay Library catalog numbers for example photographs are provided in *Supplementary Material: Manakin plumage maturation descriptions*. Plumage descriptions were drawn from literature reports, banding records, photographs, and museum specimens. For most taxa in our dataset (35/40), descriptions included information from published or unpublished banding records of recaptured individuals. For the few remaining taxa (5/40), we primarily relied on museum specimens. We excluded descriptions that relied on museum specimens alone if they implied autapomorphic characters in ways that were equivocal, confounding, or vague when compared to robust banding records of closely related species (e.g., description of *Pseudopipra pipra coracina* in Zimmer 1936). Our final dataset was missing two monotypic genera—*Protopelma* and *Ilicura* (Ohlson et al. 2013; van Els et al. 2023)—but included all presently-described variations in manakin plumage maturation. We followed the taxonomy in IOC World Bird List v15.1 (Gill et al. 2025) with two exceptions: (1) we split *Pseudopipra* into multiple species following Berv et al. (2021) and (2) we retained the genus *Antilophia*, rather than lumping it within *Chiroxiphia*.

**Figure 2.**
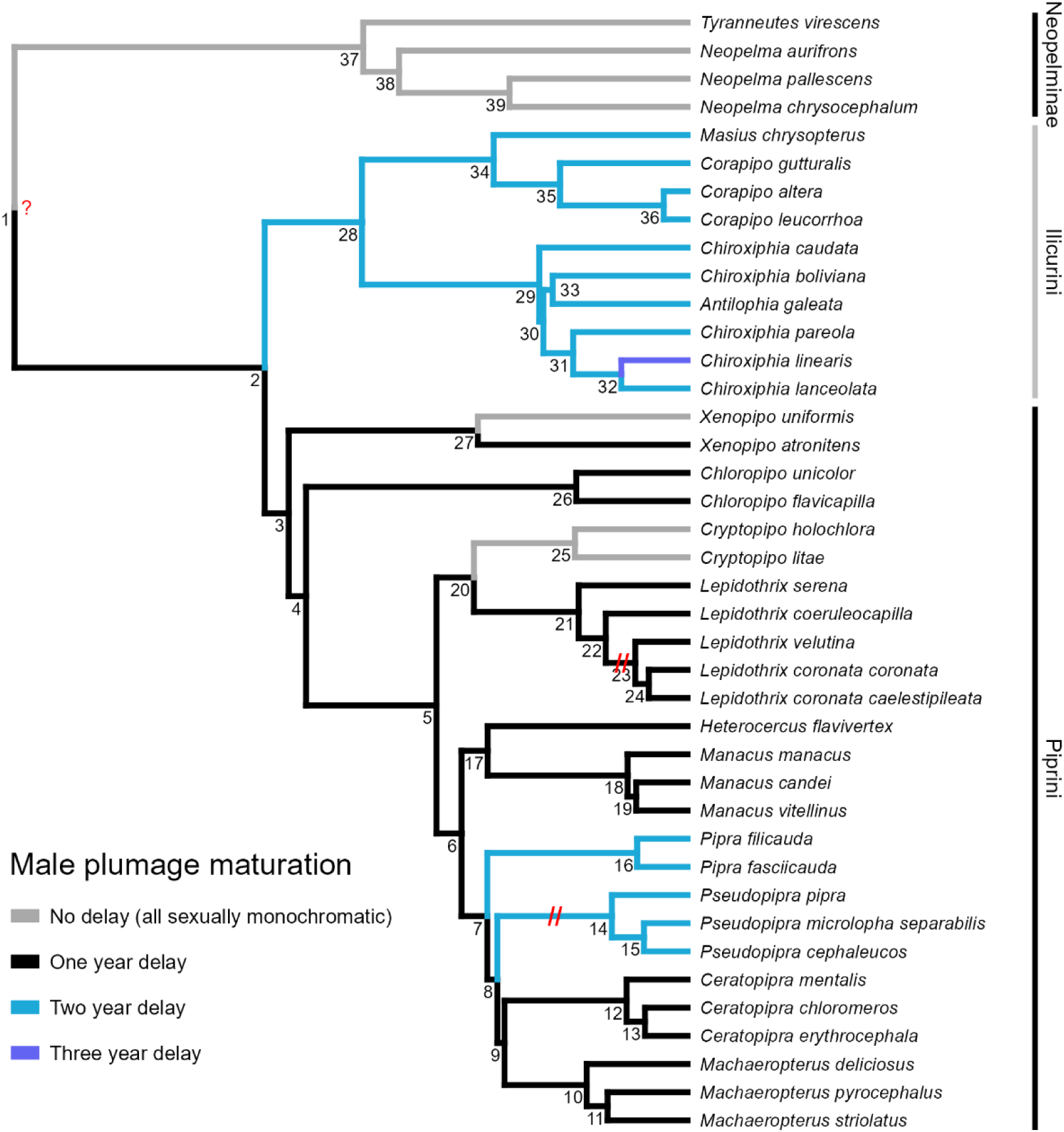
Phylogenetic history of male plumage maturation in manakins. Edges colored based on the reconstruction of developmental schedules across a set of individual plumage patch characters at each child node. Red question mark indicates uncertainty surrounding phenotypes at the root node. Node numbers correspond to Table 4.

Characters in our dataset were plumage patches (characters 1–28; Table 1). Because there are no underlying morphological structures (e.g., follicles, feather tracts, body segments) that define, circumscribe, or evolutionarily constrain plumage patches on a bird’s body, we defined a plumage patch as a module of feathers that covary in structure and color (Prum and Dyck 2003). We identified plumage patches in terms of human vision, rather than the tetrahedral color space available to avian vision (Stoddard and Prum 2008). Although some manakin plumages include ultraviolet reflectance not visible to humans (Morales-Betancourt and Castaño-Villa 2018), there is no evidence for qualitatively distinct, cryptic ultraviolet patches in these birds.

**Table 1.** Manakin plumage patch characters.

| No. | Description |
| --- | --- |
| 1 | Female body feathers |
| 2 | Female wing feathers |
| 3 | Body or back black |
| 4 | Body blue |
| 5 | Breast and/or belly brown, with or without light streaks |
| 6 | Breast, belly, and face solid yellow |
| 7 | Breast, belly, and head diffuse yellow and red |
| 8 | Belly yellow |
| 9 | Belly and rump gray |
| 10 | Flight feathers black |
| 11 | Hidden wing patch on edges of remiges and rectrices yellow |
| 12 | Hidden wing patch on secondary remiges white |
| 13 | Visible wing fringe white |
| 14 | Mask around eyes black |
| 15 | Hidden, erectile crown yellow |
| 16 | Visible crown (or forecrown) yellow/orange/red |
| 17 | Crown extended into red cape |
| 18 | Visible crown (or forecrown) white/gray/blue |
| 19 | Visible, flat cap black |
| 20 | Erectile throat white (or yellow), extending forward (or lateral) |
| 21 | Throat spot yellow |
| 22 | Scaly nape yellow/orange/red with black feather tips |
| 23 | Collar white (or yellow) wrapping around throat and nape |
| 24 | Head black |
| 25 | Head gray |
| 26 | Head yellow/orange/red |
| 27 | Back blue |
| 28 | Rump blue |

The only plumage patches in our dataset are those needed to analyze differences between predefinitive and definitive plumages within and across manakin taxa. We did not characterize juvenile plumages, which are drab, sexually monomorphic, and generally resemble female definitive plumages wherever described (Kirwan and Green 2011; Johnson and Wolfe 2018). Further, we did not score the extensive variation in female plumage among manakin taxa. Instead, we include only two general characters—female body feathers and female wing feathers (characters 1 and 2; Table 1)—applied across all taxa. Similarly, we did not analyze the diversity of male definitive plumages when that diversity was unrelated to variation in predefinitive plumages; for example, we did not code short versus long rectrices in *Chiroxiphia,* streaked versus unstreaked bellies in *Machaeropterus*, or the different colors of leg tufts in *Ceratopipra* (Kirwan and Green 2011). Additional characters would be necessary for comprehensive analyses of male and, especially, female definitive plumage evolution.

Character states in our dataset represented the developmental schedule of a plumage patch (states ϕ, A–I; Table 2). Each character state corresponds to an ordered pair of values: (*appearance age, number of molts*). The first value indicates the age at which a young male first develops the plumage patch, in months. The second value indicates the number of molts through which a plumage patch is re/generated (*n* = indefinitely, for the remaining life of the bird). State ϕ means the plumage patch is absent.

**Table 2.** Discrete character states, each of which represents the developmental schedule of a plumage patch character with an ordered pair of values: (*appearance age*, *number of molts*). Appearance age is in months. When the number of molts = *n*, the plumage patch is iteratively regenerated for the remaining life of the bird.

| Code | State |
| --- | --- |
| $\phi$ | Absent |
| A | 3, *n* |
| B | 15, *n* |
| C | 27, *n* |
| D | 3, 1 |
| E | 12, 1 |
| F | 15, 1 |
| G | 3, 2 |
| H | 15, 2 |
| I | 3, 3 |

Manakins molt out of their juvenile plumage via a partial, “preformative” molt (excluding flight feathers) at ∼3 months old, followed by complete, “prebasic” molts at ∼12-month intervals (Ryder and Durães 2005; Johnson and Wolfe 2018; Schaedler et al. 2021b). Therefore, a character state with appearance age 3 indicates the preformative molt, while appearance age 15 indicates the first complete prebasic molt, and so on (Table 2). Some *Chiroxiphia* species appear to have an additional “presupplemental” molt ∼12 months old (DuVal 2005). In our dataset, this presupplemental molt only ever generates a character for the first time (the black mask, character 14; Table 1), and never eliminates or regenerates any existing patches. In contrast, all patches are eliminated and potentially regenerated at every prebasic molt.

Each character state unambiguously represents the developmental schedule of a plumage patch (*Supplementary Material: Deriving characters from graph structures*). For example, a plumage patch with character state D = (3, 1) is generated once at the preformative molt ∼3 months old, is never regenerated, and therefore disappears with the prebasic molt ∼15 months old. In contrast, a plumage patch with character state C = (27, *n*) is first generated at the prebasic molt ∼27 months old and then regenerated annually, at every subsequent prebasic molt, for the remaining life of the bird. When combined, our 28 plumage patch characters (Table 1) and 10 developmental schedule states (Table 2) can unambiguously represent the male plumage maturation process for each taxon in our dataset (Table 3). For example, *Xenopipo atronitens* is coded:

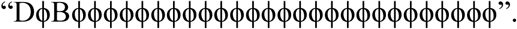

**Table 3.**
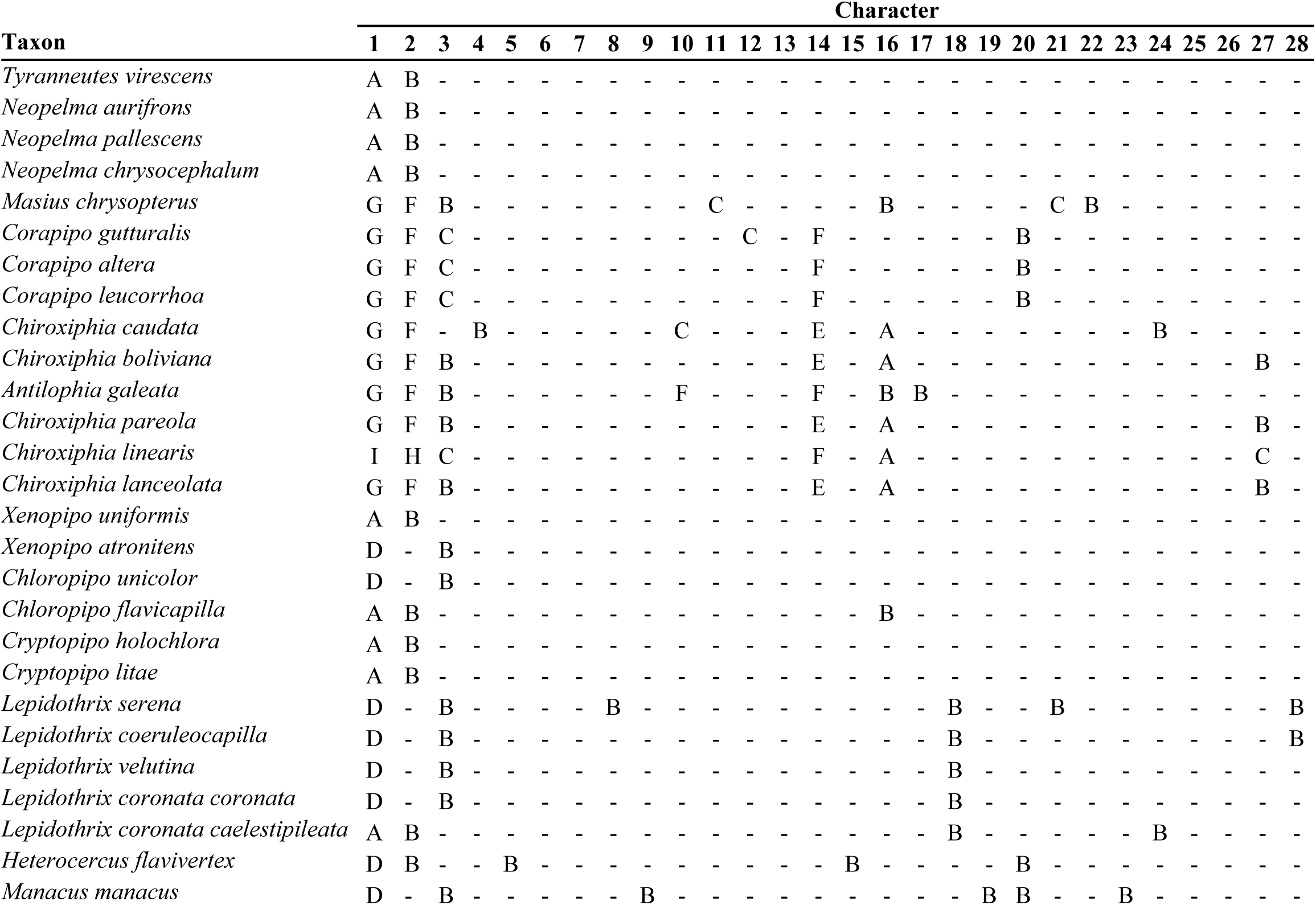
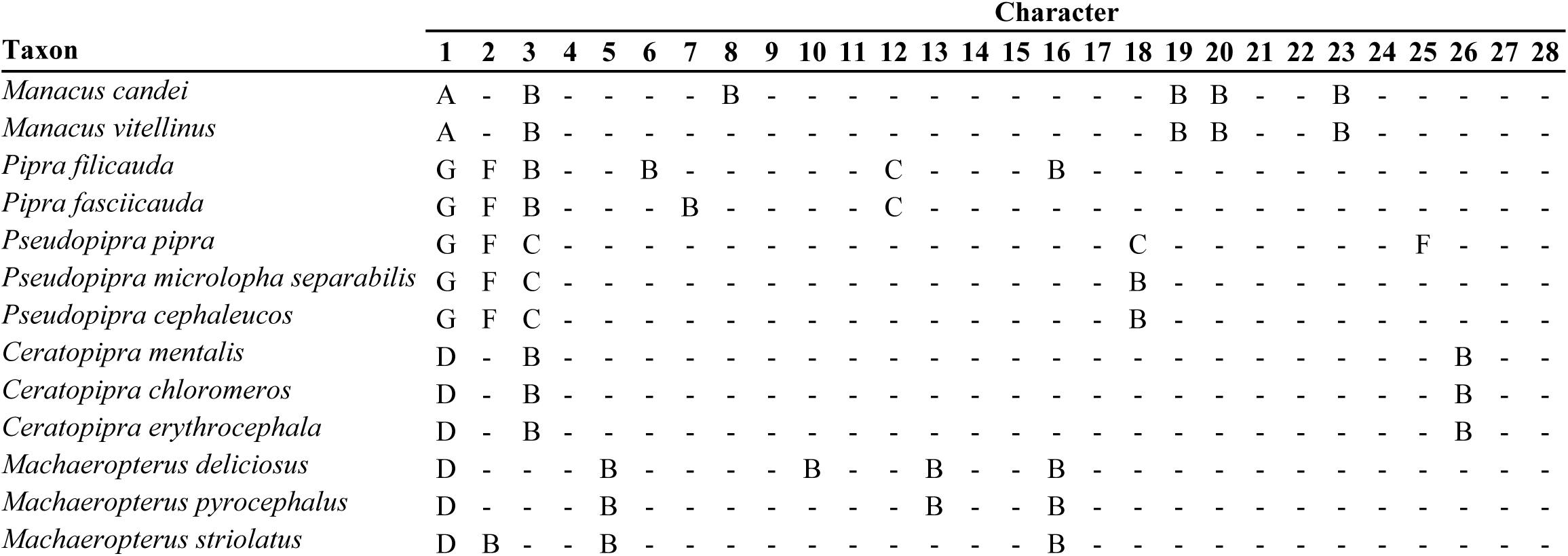
Comparative dataset for male plumage maturation in 40 manakin taxa. Absent characters (state ϕ) shown with a dash.

In *X. atronitens*, male plumage maturation includes female contour feathers at the preformative molt—character 1, state D = (3, 1)—followed by a prebasic molt into a black body plumage, which is then regenerated at each prebasic molt for the remaining life of the bird— character 3, state B = (15, *n*).

In contrast, *X. uniformis* is coded:

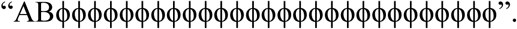

In *X. uniformis*, male plumage maturation includes definitive female contour feathers at the preformative molt—character 1, state A = (3, *n*)—and definitive female flight feathers in the subsequent prebasic molt—character 2, state B = (15, *n*). Both patches are regenerated via prebasic molts for the remaining life of the bird. In other words, *X. atronitens* spends one year in a female plumage followed by a black, sexually dichromatic male definitive plumage, whereas *X. uniformis* is sexually monochromatic with no DPM.

### Ancestral state modeling

For phylogenetic analysis, we used the time-calibrated, maximum likelihood (highly filtered UCE “HGAPF” dataset) suboscine tree from Harvey et al. (2020), pruned to include only taxa in our dataset (Fig. 2). The Harvey tree lacked some taxa for which we had plumage maturation data, so we grafted branches for *Lepidothrix* (Moncrieff et al. 2022) and *Pseudopipra* (Berv et al. 2021). Because these trees were not ultrametric, we calibrated their divergence times with the ape::chronos function in R v. 4.4.1 (Paradis and Schliep 2019; R Core Team 2024). For *Lepidothrix*, the common ancestor of *L. coeruleocapilla* and *L. coronata* was present in both the Harvey and Moncrieff trees. Thus, we used the age of that ancestor in the Harvey tree (1.84 mya) to calibrate the Moncrieff *Lepidothrix* branches. For *Pseudopipra*, which is represented by only one tip in the Harvey tree, we used an estimated age of 2.437 mya for the common ancestor of *Pseudopipra* (Castro-Astor 2014). The resulting divergence times are rough estimates, especially given the different genetic datasets used for different tips in the Berv *Pseudopipra* tree.

To reflect ongoing uncertainty in the manakin phylogeny, we replicated our analysis on the maximum likelihood (concatenated 95% UCE dataset) tree from Leite et al. (2021). Compared to the Harvey tree (Fig. 2), the Leite tree (Fig. S1) shows different species relationships within *Chiroxiphia* and resolves *Pseudopipra* (rather than *Machaeropterus*) as sister to *Ceratopipra*. The Leite tree is not ultrametric and lacked several species from our dataset. For this alternative analysis, we simply analyzed the 34 taxa from our dataset that were featured in the Leite tree. All results using the Leite tree were consistent with our main analysis using the Harvey tree (*Supplementary Material: Alternative phylogeny*).

We modeled ancestral states separately for each character using maximum likelihood estimation in the ape::ace function. We tested four continuous-time, discrete state Markov transition models (Pagel 1994) for each character: equal rates (ER), symmetric rates (SYM), all rates different (ARD), and a custom model (ERϕ). The ERϕ model differed from ER by providing distinct, asymmetric parameters for transitions to or from the absent state (ϕ). In other words, the ERϕ model distinguishes losing or gaining a plumage patch (e.g., state ϕ → A) from all transitions in developmental schedule, including transitions in appearance time (e.g., state A → B), transitions in number of molts (e.g., state A → D), and joint transitions in appearance time and number of molts (e.g., state B → D). Some models were equivalent for characters with a limited number of observed states (e.g., ER and SYM models are equivalent for a character with only two states). Examples of the four models are given in Table S2.

We selected the transition model with the lowest AIC score for each character (Fig. 3). Estimated transition rates from multi-parameter models were spuriously high for characters present in few tips, so we restricted characters present in <3 taxa to ER models only. Data and trees were wrangled and visualized using the *tidyverse* and *ggtree* packages in R (Yu et al. 2017; Wickham et al. 2019).

**Figure 3.**
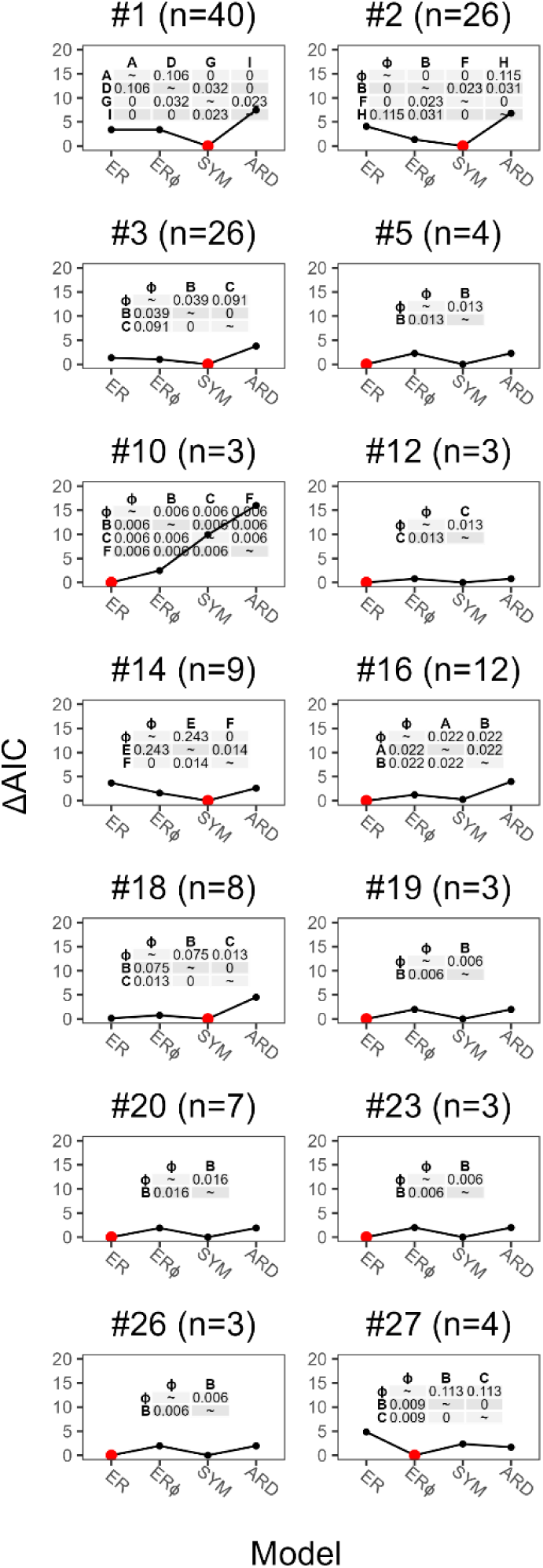
Character transition models for 14 plumage patch characters that appeared in >3 taxa. Panels are labeled by character number (n = number of tips where the character is present). Four discrete transition models were tested for each character: equal rates (ER); equal rates except for asymmetric transitions to/from the absent state (ERϕ); symmetric rates (SYM); all rates different (ARD). Red points indicate the preferred model for each character, which had the lowest Akaike Information Criterion (AIC) score. Inset tables show maximum likelihood transition rate parameters under each character’s preferred model. The remaining 14 characters, not shown, were present in <3 taxa and restricted to an ER model with uninformative transition rates.

## Results

Our phylogenetic analysis revealed the evolutionary origins, elaborations, and losses of male DPM in manakins (Fig. 2). We modeled the evolution of separate plumage patch characters (Fig. 4), each under their own preferred transition model (Fig. 3). Fourteen of 28 characters (50%) were present in only one or two taxa, meaning they were restricted to an ER model. Ten of those 14 rare characters had a single transition from absent (ϕ) to a single present state (Fig. S2). The other four rare characters (characters 8, 13, 24, 28) had two independent transitions from absent (ϕ) to a single present state (Fig. S2). For the remaining 14 characters—all present in three or more taxa—eight used ER models, five used SYM models, and one (character 27) used the custom ERϕ model (Fig. 3). Note the uninformative transition rates in characters that originate in a single state with no subsequent evolution (e.g., characters 5 and 24; Figs. 3, S2).

**Figure 4.**
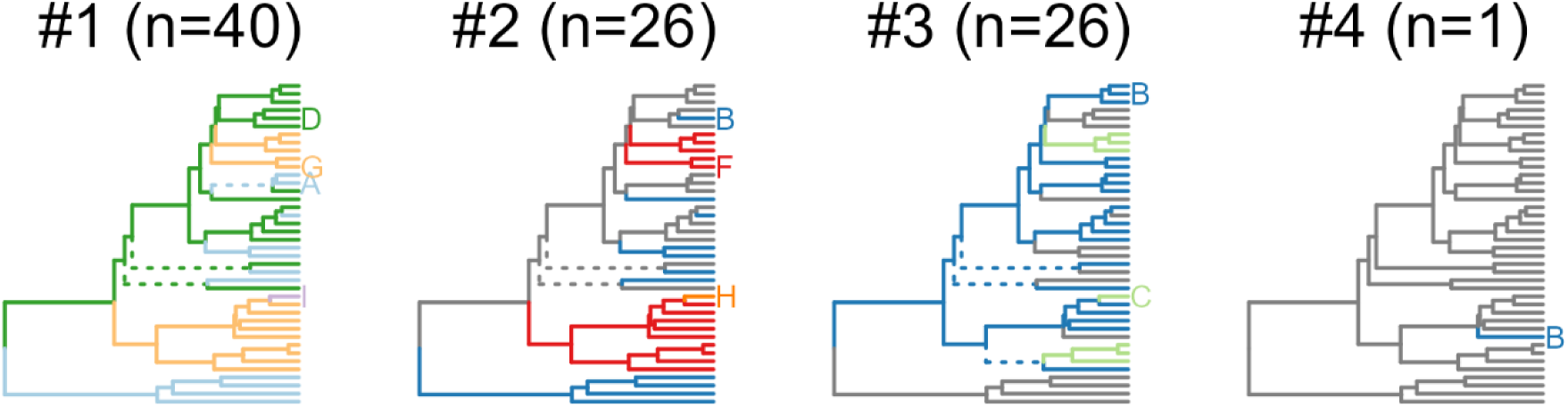
Example of plumage patch character histories (characters 1–4). Panels are labeled by character number (n = number of tips where the character is present). Edges colored according to the maximum likelihood character state at each child node. Dashed lines indicate uncertainty (scaled likelihood <0.75). See Figure S2 for additional characters.

As with crown taxa (Table 3), concatenating the 28 plumage patch characters in their maximum likelihood ancestral states (Fig. 4) allowed us to reconstruct the male plumage maturation process for each common ancestor in the phylogeny (Table 4, with node numbers corresponding to Fig. 2). Below, we walk through the inferred evolutionary history of DPM in manakins.

**Table 4.**
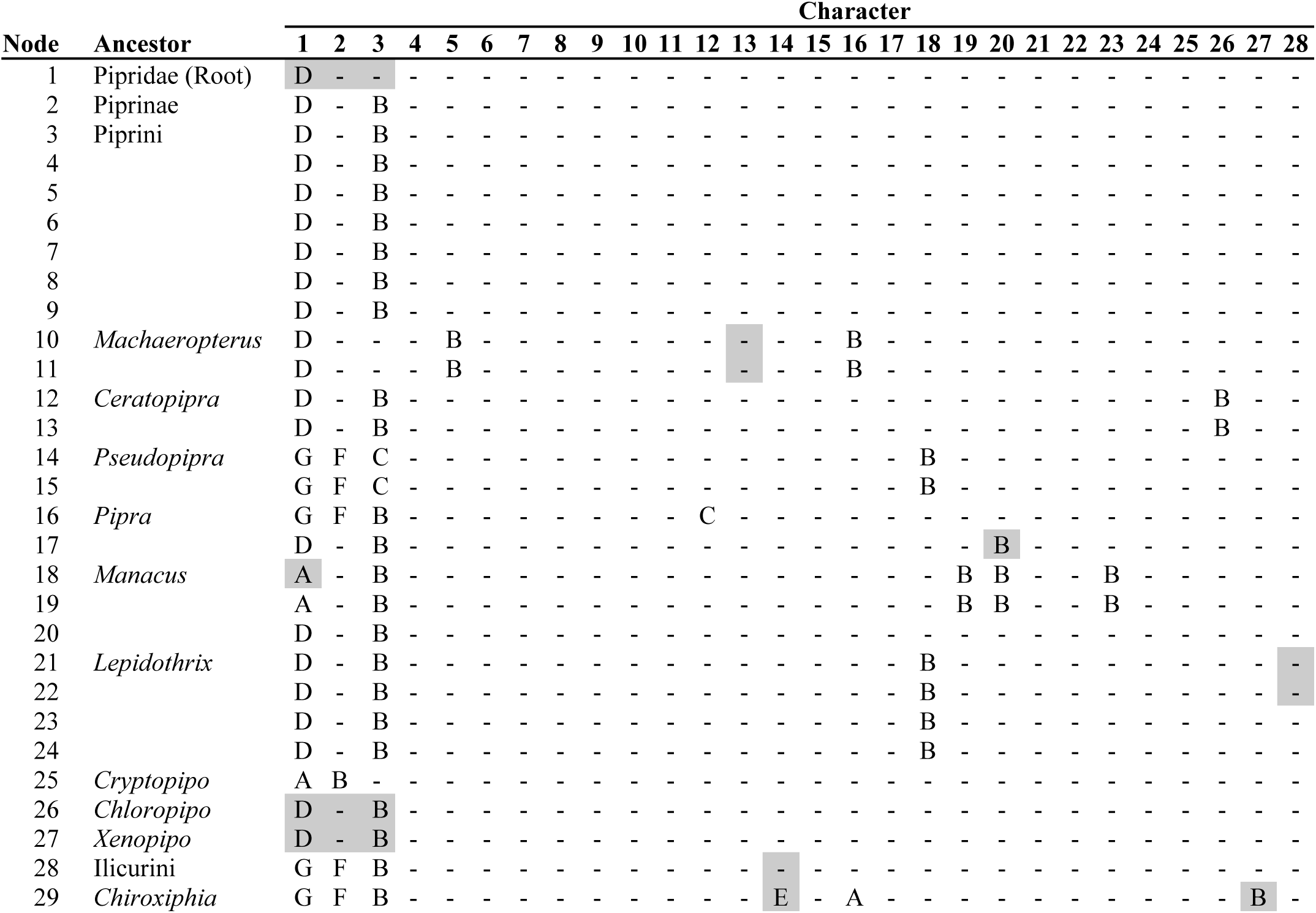
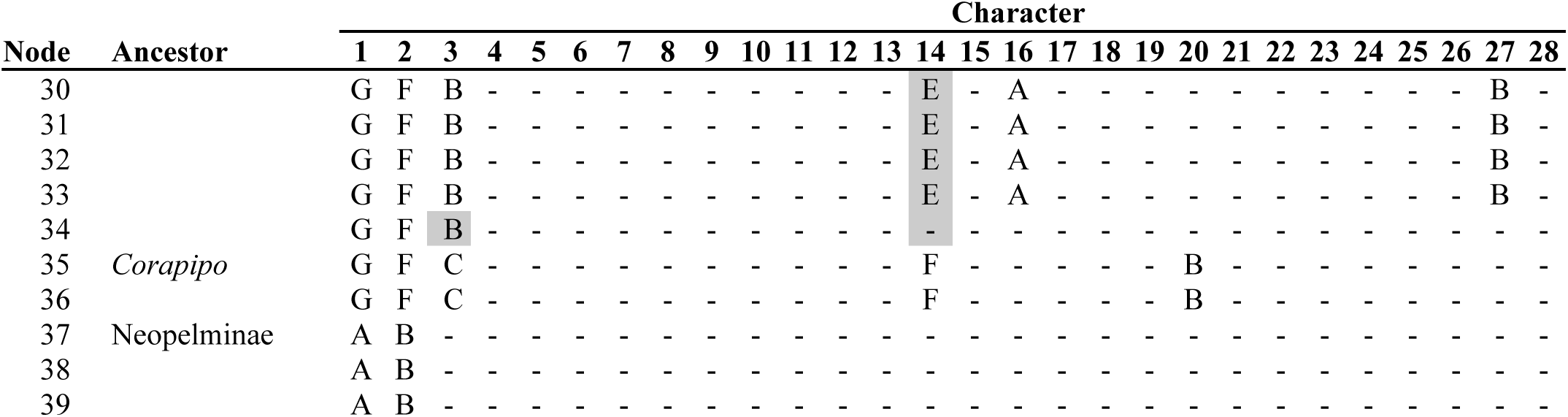
Ancestral male plumage maturation for manakins. Maximum likelihood character states shown for each of 28 plumage patch characters. Uncertain states indicated in gray (scaled likelihood <0.75). All states had scaled likelihood >0.50. Absent characters (state ϕ) shown with a dash. Node numbers correspond to Figure 2.

### Neopelminae

The Neopelminae, or tyrant-manakins, are the sister group to all other manakins (Fig. 2). All Neopelminae are sexually monochromatic with no DPM (though see notes on *Tyranneutes virescens* in *Supplementary Material: Manakin plumage maturation descriptions*). Young males in this lineage obtain female body feathers in the preformative molt, female flight feathers in the subsequent prebasic molt, and thereafter regenerate that plumage for the rest of their lives (Table 3). Although our dataset is missing *Protopelma chrysolophum* and three other poorly-studied Neopelminae (all of which are also sexually monochromatic), we resolved the ancestor of Neopelminae as sexually monochromatic with no DPM (node 37).

### Piprinae

The Piprinae include all other manakins, in tribes Ilicurini and Piprini (Fig. 2; Ohlson et al. 2013). In all sexually dichromatic Piprinae, males have at least one year of DPM (Table 3). Accordingly, we resolved the origin of one-year DPM at the common ancestor of Piprinae (node 2; Table 4). We were unable to resolve a specific plumage maturation process for the common ancestor of Pipridae (node 1) due to the high diversity of male plumage elements across crown taxa and the primary divergence between the sexually monochromatic Neopelminae and the sexually dichromatic Piprinae (Fig. 2).

Males in the Piprinae ancestor (node 2) spent one year in a fully female predefinitive plumage, before passing into a sexually dichromatic male definitive plumage with black body feathers, via prebasic molt at ∼15 months old (character 3, state B). Most Piprini manakins in our dataset retained this ancestral DPM process, including *Xenopipo atronitens*, *Chloropipo*, *Lepidothrix*, *Heterocercus*, *Manacus*, *Ceratopipra*, and *Machaeropterus*. Young males in these taxa spend one year in a fully female predefinitive plumage before molting into lineage-specific male definitive plumages (Table 3).

Following the origin of one-year DPM in Piprinae, there were three independent origins of prolonged, two-year DPM (Fig. 2). We placed these origins in the respective ancestors of: Ilicurini (node 28), *Pipra* (node 16), and *Pseudopipra* (node 14). All three lineages evolved to not only prolong DPM but also exhibit lineage-specific signals of male status within new predefinitive plumage stages.

### Ilicurini

The Ilicurini include genera *Ilicura* (no data), *Masius*, *Corapipo, Chiroxiphia*, and *Antilophia* (Fig. 2). All Ilicurini manakins in our dataset have two-year DPM except for *C. linearis*, which extends development into three-year DPM (Fig. 2). Male plumage maturation in the Ilicurini ancestor (node 28) featured one year in female body feathers (character 1, state G), followed by one year in a mix of regenerated female body feathers, newly molted female flight feathers (character 2, state F), and some sexually dichromatic black body feathers (character 3, state B). This ancestor then obtained a black, male definitive plumage via prebasic molt ∼27 months old. There was uncertainty surrounding the appearance of a transient black mask (character 14) earlier in ontogeny, with scaled likelihoods of 56.4% for the absent state, 35.8% for state F (appearing once via prebasic molt ∼15 months old), and 7.7% for state E (appearing once via presupplemental molt ∼12 months old).

Within each Ilicurini lineage, predefinitive plumages continued to evolve to maintain lineage-specific signals of male identity. In *Masius chrysopterus* (Fig. 1A), males pass from the first, female predefinitive plumage into a second predefinitive plumage with black body feathers (character 3, state B), a yellow forecrown (character 16, state B), and scaly nape (character 22, state B) via prebasic molt ∼15 months old (Fig 1A; Table 3). All *Corapipo* manakins (descending from node 35) pass from the first, female plumage to a second predefinitive plumage with a black mask (character 14, state F) and white throat (character 20, state B; Fig. 1B).

Species in *Chiroxiphia* (Fig. 1C) and *Antilophia* (Fig. 1D) elaborated their predefinitive plumages in two ways. First, *Chiroxiphia* evolved to reveal male status via preformative molt ∼3 months, rather than spending a year in a fully female plumage. All *Chiroxiphia* taxa obtain a red crown at the preformative molt (character 16, state A), a feature resolved for their common ancestor (node 29). For *A. galeata*, in contrast, the appearance of red forecrown feathers is delayed until ∼15 months (character 16, state B), when it appears in conjunction with some black body feathers (character 3, state B) and parts of the red cape (character 17, state B).

The second kind of evolutionary elaboration within *Chiroxiphia* occurred in *C. linearis*. Unique among manakins, *C. linearis* evolved three-year DPM (Fig. 2, Table 3). The common ancestor of *Chiroxiphia* obtained a red crown via preformative molt ∼3 months old, plus transient black mask via presupplemental molt ∼12 months, plus a blue mantle and additional black body feathers via prebasic molt ∼15 months (nodes 29–33). Male definitive plumage is thus obtained via the next prebasic molt ∼27 months. This condition is apparently retained in *C. boliviana*, *pareola*, and *lanceolata*. In contrast, *C. linearis* evolved to pass through the same predefinitive stages at a slower rate: red crown via preformative molt ∼3 months old, plus transient black mask via prebasic molt ∼15 months, plus a blue mantle and additional black body feathers via prebasic molt ∼27 months (Table 3). The definitive plumage in *C. linearis* is thus obtained ∼39 months.

### Piprini

Moving on from Ilicurini, there were two additional origins of two-year DPM in Piprini manakins. One was in the ancestor of *Pipra* (node 16). The two *Pipra* species in our dataset (*P. filicauda* and *P. fasciicauda*) share a developmental schedule with two predefinitive plumage stages. However, each species evolved to exhibit autapomorphic, male-like carotenoid plumage patches in the second predefinitive plumage, via prebasic molt ∼15 months (characters 6 and 16 in *P. filicauda*, character 7 in *P. fasciicauda*). Note that because lineage-specific carotenoid plumage patches are coded differently in the two *Pipra* crown taxa, the common ancestor of *Pipra* was inferred to have black plumage elements (character 3) and a definitive white wing patch (character 12), but artifactually lacks carotenoid plumage patches.

The other origin of two-year DPM was in the ancestor of *Pseudopipra* (node 14). In all *Pseudopipra*, the male definitive plumage is black with a white crown. In *P. pipra,* females are largely green, and males obtain a transient, predefinitive gray head via prebasic molt ∼15 months (character 25, state F; Fig. 1E), waiting until the next prebasic molt ∼27 months to obtain the black body (character 3, state C) and white crown (character 18, state C). In the *P. cephaleucos* and *P. microlopha separabilis* clade, females are green with a gray head, and young males obtain a white crown (*P. cephaleucos*) or a unique gray crown (*P. microlopha separabilis*; both coded character 18, state B) beginning ∼15 months, before obtaining male definitive plumage ∼27 months.

### Loss of DPM

Results indicated two losses of DPM in the manakin phylogeny (Fig. 2). One loss occurred in *Xenopipo uniformis* (Table 3), and the other occurred in the common ancestor of *Cryptopipo* (node 25; Table 4). These losses are coincident with the secondary evolution of sexually monochromatic definitive plumage.

## Discussion

Our phylogenetic analysis resolved the key events in the evolution of male plumage maturation in manakins (Fig. 2). All Piprinae manakins (i.e., all manakins excluding the sexually monochromatic tyrant-manakins, or Neopelminae) descend from a common ancestor in which young males spent one year in a female plumage before developing a sexually dichromatic, male definitive plumage. Some lineages have retained this condition, including *Xenopipo atronitens*, *Chloropipo, Lepidothrix*, *Heterocercus*, *Manacus*, *Ceratopipra*, and *Machaeropterus*. Two lineages with secondarily derived sexual monochromatism—*Xenopipo uniformis* and *Cryptopipo*—have lost male DPM, insofar as they have lost all distinctions between female and male plumage. Three lineages—Ilicurini, *Pipra*, and *Pseudopipra—*evolved to have longer, two-year DPM. Within Ilicurini, *Chiroxiphia linearis* evolved even longer, three-year DPM. Each case of prolonged DPM evolved to reveal signals of young male status during later predefinitive plumage stages, and each continued evolving to maintain those signaling properties whenever signals of maleness themselves diversified (Tables 3, 5).

### Social signaling

The evolutionary history of manakin DPM provides comparative evidence for a social signaling function (McDonald 1993a; Schaedler et al. 2021b), as opposed to alternate functions such as crypsis or female mimicry (Rohwer 1978; Rohwer et al. 1980; Hawkins et al. 2012). The three independent origins of two-year DPM (Ilicurini, *Pipra*, and *Pseudopipra*) and the single origin of three-year DPM (*C. linearis*) all revealed developmental periods during which young males have plumages that signal their status as young males. For example, the second predefinitive plumage stage (∼15–27 months) in *Corapipo* features a white throat and black mask, and the second stage in *Pipra* features yellow or red carotenoid patches (Fig. 1; Tables 3, 5). The evolution of predefinitive plumages that clearly indicate male status is consistent only with a social signaling function. This evolutionary pattern runs counter to crypsis and female mimicry hypotheses, which predict that young males evolve DPM only insofar as they might prolong the time spent in drab (crypsis) or female (mimicry) plumages.

Wherever prolonged DPM evolved and aspects of male signaling diversified, predefinitive plumages codiversified alongside them. All Ilicurini manakins descend from a common ancestor with prolonged, two-year DPM (Fig. 2). Thus, the second predefinitive plumage stage is homologous across Ilicurini. Yet Ilicurini manakins have evolved lineage-specific signals of male identity at that homologous stage. These signals are themselves non-homologous among lineages. For example, in their definitive male plumages, *Masius* exhibits a yellow forecrown, *Corapipo* a white throat, and *Chiroxiphia* a red crown. In parallel, the second predefinitive plumage stage in *Masius* includes a yellow forecrown, in *Corapipo* a white throat, and in *Chiroxiphia* a red crown, among other things (Fig. 1). Even though the developmental stages of DPM are homologous, the specific plumage patches molted in are not. Rather, these predefinitive plumage patches track the apomorphic features of male definitive plumages in each lineage, maintaining their potential to function as social signals of both sex and developmental stage. To our knowledge, this is a novel finding for any avian clade.

The evolution of predefinitive plumages in *Pseudopipra* supports a social signaling function in a different, unique, and unexpected way. Male definitive plumages are largely indistinguishable across *Pseudopipra*, all featuring a black body with a white crown. Despite stasis in male definitive plumages, *Pseudopipra* shows diversification in both female definitive and male predefinitive plumages. In the second male predefinitive plumage stage, *P. pipra* exhibits a gray head (Fig. 1E), whereas *P. cephaleucos* and *P. microlopha separabilis* exhibit a gray head plus a white (or gray) crown (Zimmer 1936; Johnson and Wolfe 2018; Berv et al. 2021). Although we did not explicitly analyze the evolution of female plumages, our observations and previous descriptions (Kirwan and Green 2011; Johnson and Wolfe 2018) indicate that female plumage in *P. pipra* is overall green, whereas female plumage in *P. cephaleucos* and *P. microlopha separabilis* is green with a gray head. In other words, a gray head is sufficient to distinguish young males from females in *P. pipra*, but not in *P. cephaleucos* and *P. microlopha separabilis*. The latter lineages are the ones to evolve the white (or gray) crown patches in the second predefinitive plumage.

Thus, in *Pseudopipra*, it appears that predefinitive plumages evolved not through codiversification with maleness, but rather through counter-diversification with femaleness. The process is different, but the result is the same: a plumage that indicates young male status. One important caveat is that our dataset includes just three *Pseudopipra* taxa out of potentially 10 or more species (Berv et al. 2021). Fully understanding the evolution of *Pseudopipra* plumages will require a more complete analysis of the genus, including explicit attention to female plumage diversity and descriptions of Andean taxa that diverged prior to the *Pseudopipra* taxa in our dataset. Studies of *Pseudopipra* will be particularly interesting because the genus includes predefinitive plumage patches that differ from both male and female definitive plumages, specifically the gray crown in young male *P. microlopha separabilis* (Zimmer 1936).

Our phylogenetic analysis alone cannot provide evidence for a social signaling function at the primary origin of DPM in the ancestor of Piprinae manakins (Fig. 2). In this ancestor and several sexually dichromatic descendants, young males spend one year in an unornamented female plumage with no patches to signal maleness. However, as Schaedler et al. (2021b) remark, one may still reject a female mimicry function for these predefinitive plumages in Piprinae insofar as young males signal their status with social and sexual behaviors. We also mark an exception for *Chiroxiphia*, in which young males evolved to exhibit a feature of male identity—the red crown—as early as the preformative molt ∼3 months old (Tables 3, 5; DuVal 2005; Doucet et al. 2007a; Mallet-Rodrigues and Dutra 2012).

In general, plumage evolution in manakins is too dynamic to accurately reconstruct phenotypes at the root of the family. The clade is characterized by an explosive phenotypic diversity that makes it challenging to resolve ancestral conditions (Prum 1997). Furthermore, the initial divergence in Pipridae is between two radically different clades: the sexually monochromatic Neopelminae and sexually dichromatic Piprinae. Lastly, it is difficult to polarize character changes with an outgroup because Pipridae is sister to the rest of Tyrannides (Harvey et al. 2020). This group encompasses several large lineages, including hundreds of tyrant flycatchers (Tyrannidae) and dozens of cotingas (Cotingidae), the latter of which especially features its own diversity in sexual dichromatism and DPM (Kirwan and Green 2011).

### Sociosexual development

We conclude that phylogenetic patterns in plumage development in three manakin clades –Ilicurini, *Pipra*, and *Pseudopipra*–independently support the social signaling hypothesis for DPM. Intraspecific field studies, especially in *Chiroxiphia*, demonstrate how predefinitive social signals may benefit young males entering display leks (Foster 1987; McDonald 1993a; DuVal 2013; Schaedler et al. 2021a). For example, in Long-tailed Manakins (*Chiroxiphia linearis*), young males enter a complex, hierarchical competition with unrelated males at display sites, often perform as a secondary “beta” dancer in cooperative male-male displays, eventually graduate to an “alpha” position, and subsequently refine their own display with male partners (Trainer et al. 2002; McDonald 2010). This process can take each bird more than a decade (McDonald 1993b). Male-male relationships in *Chiroxiphia* are especially elaborate given their cooperative, coordinated sexual displays (Foster 1977). In *Pipra filicauda*, males only occasionally perform coordinated displays, but they must still maintain their display sites by developing status among male coalitions (Ryder et al. 2008, 2011; Vernasco et al. 2020). In either case, predefinitive plumages that explicitly signal young male status may reduce the risk of conflict while young birds establish durable male-male relationships (McDonald 1993a).

Future studies are needed to investigate the content of male sociosexual development and male-male relationships across a wider range of manakins, especially in lineages with simpler displays such as *Pseudopipra* (Berv et al. 2021). Additional work can also investigate more direct ties between sociosexual development and DPM. Are there consistent differences in male sociosexual development between those lineages with simple, one-year DPM (e.g., *Ceratopipra*) and those with more elaborate predefinitive plumages (e.g., some *Pseudopipra*)? Is the evolution of DPM merely a consequence of risks inherent to male-male relationships at the lek, or does the evolution of predefinitive plumages expose new possibilities for building those relationships? For example, we note that prolonged delayed plumage maturation evolved in Ilicurini prior to the transition from occasional multi-male displays (as in e.g., *Masius chrysopterus*) to fully cooperative, multi-male displays performed for a female audience (as in *Chiroxiphia*; Prum 1994).

### Developmental evolution

Beyond social signaling, our results shed light on the developmental evolution of manakin plumages. Specifically, we resolved two novel phenotypes that evolved through heterochronic shifts (Gould 1977). The first involves the origins of sexual monochromatism in *Xenopipo uniformis* and *Cryptopipo.* Ribeiro et al. (2015) show that both lineages likely evolved sexual monochromatism from sexually dichromatic ancestors. Our analysis further demonstrates that these ancestors had DPM, with young males spending one year in a green, female plumage (Fig. 2). From this perspective, the evolution of sexual monochromatism involves not merely a loss of male definitive plumage, but rather the indefinite regeneration of the female plumage. In other words, sexual monochromatism in *Cryptopipo* and *Xenopipo uniformis* evolved through the suspension of ancestral male plumage maturation, corresponding to Gould’s (1977: 229) concept of “progenesis.” The deep developmental mediation of sexual dichromatism in manakins is further supported by the fact that older females in several species obtain those features of male plumage that appear in the youngest males (Graves 1981; see e.g., detailed examination of *C. linearis* crowns in Doucet et al. 2007a). The idea that heterochronic shifts underlie evolutionary diversity in avian dichromatism was clearly discussed by Björklund (1991).

A second heterochronic shift involves the origin of extended, three-year DPM in *C. linearis*. Young *C. linearis* males pass through three predefinitive plumage stages: (A) red crown ∼3–15 months, (B) black mask ∼15–27 months, and (C) blue back ∼27–39 months. Each stage lasts one year in *C. linearis*, meaning males achieve a definitive plumage ∼39 months old (Doucet et al. 2007a). Our analysis suggests *C. linearis* descended from an ancestor with the same three predefinitive stages, but condensed timing in the first two stages: (A) red crown ∼3– 12 months, (B) black mask ∼12–15 months, (C) blue back ∼12–27 months. This ancestral schedule, retained in e.g., *C. lanceolata*, results in definitive plumage ∼27 months old (DuVal 2005). The three-year plumage maturation in *C. linearis* evolved through “neoteny” (Gould 1977 p. 229), in which young males pass through ancestral developmental stages but at a slower rate. Because plumage maturation in *C. linearis* is both slower *and* longer, it does not result in either paedomorphosis (i.e. the retention of an ancestral immature state in adult morphology) or hypermorphosis (i.e., the development of novel morphology beyond that achieved in ancestral adults).

### Iterative developmental homologies in a phylogenetic context

Most broadly, the history of manakin plumages highlights new levels of complexity in the study of developmental evolution. This system involves at least four tiers of homology, all of which describe relationships between components of dynamic, developmental processes rather than static, adult phenotypes (Laubichler 2000; Gilbert and Bolker 2001; DiFrisco and Jaeger 2021). First, there is the serial homology between plumages as they are iteratively regenerated on the body of an individual bird (e.g., the plumage in the first, second, and *n*th year of life in an individual White-throated Manakin, *Corapipo gutturalis*). Extending previous definitions of serial homology (Owen 1848; DiFrisco et al. 2023), a plumage is an iteration of a single character across ontogenetic time, rather than an iteration across morphological space. Second, there is special homology between equivalent developmental stages across taxa, or discrete ontogenetic homology (e.g., the correspondence between the first predefinitive, second predefinitive, and definitive plumage stages in *C. gutturalis* and *M. chrysopterus,* though not their distinct, non-homologous plumage patches). Third, there is special homology of developmental schedule, or continuous ontogenetic homology (e.g., the shared schedule, in months, through which *C. gutturalis* and *M. chrysopterus* pass through plumage stages). Fourth, there is special homology of constituent features within each stage of the ontogenetic process (e.g., the red cap in the first predefinitive plumage shared across *Chiroxiphia*). When applied to the “adult” stage of development, this fourth level offers a traditional statement about special homology (e.g., the black-and-white male definitive plumage across *Pseudopipra*).

Each level of developmental homology also defines a level of developmental innovation or novelty (McKenna et al. 2021). For example, Piprinae manakins innovated on the temporal serial homology of plumages by differentiating an initial, green predefinitive plumage from subsequent male definitive plumages. Ilicurini manakins modified an ontogenetic process in a discrete sense by evolving a novel, second predefinitive plumage stage. *Chiroxiphia linearis* modified an ontogenetic process in a continuous sense by extending the time spent in an ancestral predefinitive plumage stage (i.e., heterochronic evolution; Gould 1977). Many lineages of manakins modified the constituent features within shared ontogenetic stages by evolving new predefinitive and definitive plumage patches. The differences between these levels of homology and innovation are obscured when one views development as a simple, continuous process. They are erased completely when one views ontogeny as mere evidence for homologies among adult phenotypes.

The complex, hierarchical nature of plumage patches (Prum and Dyck 2003) and the evolutionary lability of avian molt (e.g., Wolfe et al. 2021) makes plumage maturation an especially complicated case of developmental evolution. However, the multiple levels of developmental homology outlined here may apply to a broad range of discrete, iterative organismal structures, from instars (Truman 2019) and antlers (Clements et al. 2010) to flowers (Mason 1957). Looking forward, comparative analyses must extend methods (e.g., Jeffery et al. 2005; Tarasov 2023; *Supplementary Material: Deriving characters from graph structures*) that leave room for development to operate as not only the source, but also the substance, of evolution.

## Acknowledgements

We are grateful to many manakin researchers for sharing banding data and other observations of young manakins, including Bette Loiselle, Brandt Ryder, Dusti Becker, David McDonald, John Blake, Laura Schaedler, Lia Kajiki, Mariana Villegas, and Nicole Büttner. We thank Kristof Zyskowski and Paul Sweet for support with museum collections. Photographs were reproduced with permission from The Macaulay Library at the Cornell Lab of Ornithology. Daniel Stadtmauer, Martha Muñoz, Casey Dunn, Stephen Stearns, and Josef Uyeda provided valuable feedback on the manuscript. This study was supported by an NSF GRFP (#DGE1752134) and an American Society of Naturalists student research award for L.U.T., and the W.R. Coe Fund from Yale University.

