## Supplementary Material for "Developmental evolution of social signals in the delayed plumage maturation of manakins (Aves: Pipridae)"

**Table of Contents**

| pp. | 2 – 8 | Deriving characters from graph structures |
| --- | --- | --- |
| pp. | 9 – 52 | Manakin plumage maturation descriptions |
| pp. | 53 – 56 | Alternative phylogeny (including Fig. S1, Table S1) |
| p. | 57 | Table S2 |
| p. | 58 | Figure S2 |
| pp. | 59 – 62 | References for Supplementary Material |

**Deriving characters from graph structures**

Phylogenetic comparative methods have offered an increasingly complex set of tools to analyze discrete developmental evolution. One set of discrete methods assumes that lineages share a set of ontogenetic characters and investigates changes in the developmental relationships (i.e., relative timing) of those shared characters (Bininda-Emonds et al. 2002; Jeffery et al. 2005). Another set of discrete methods assumes that lineages share a set of developmental relationships (i.e., morphological or regulatory dependencies) and investigates changes in the form of ontogenetic characters (Tarasov 2019; Porto et al. 2024).

As described in the main text, avian plumage maturation is a discrete and iterative developmental process that can feature evolutionary changes to both the relative timing of plumage patches, and the presence/form of those patches themselves. As a result, one cannot use a comparative dataset that holds constant either a set of characters or their developmental relationships. The challenge is defining characters and character states that represent shifting biological relationships among shifting biological characters. A general solution is to define a system of character coding that represents changes to a *graph structure* (i.e., nodes, as biological characters, and edges, as relationships among those characters).

The characters and character states used in our analysis are minimum representations of the directional graphs that describe the relationships between plumages and molts during plumage maturation in manakins. As an example, consider a manakin with plumage patches P_1_, P_2_, and P_3_, which are variously generated via molts M_F_, M_S_, and M_B_ (the preformative, presupplemental, and prebasic molts, respectively). Young males in this hypothetical species gain patches P_1_ and P_2_ at 3 months (via M_F_), then gain a transient predefinitive patch P_3_ at 12 months (via M_S_), and then retain only patch P_2_ for the definitive plumage (via M_B_ at annual intervals starting at 15 months).

We can represent this plumage maturation process as a graph that features nodes for both plumage patches (P_1_, P_2_, and P_3_) and molts (M_F_, M_S_, and M_B_). Edges leading from plumage-nodes to molt-nodes represent feathers falling out, whereas edges leading from molt-nodes to plumage-nodes represent feathers growing in.

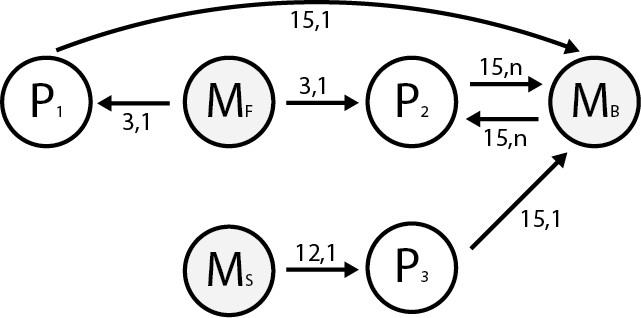
*Graph 1*

Each edge has a weight, which is vector with two elements. The value at the first index represents the first time an edge operates (in months). The value at the second index represents the number of times the edge operates (where *n* is indefinitely). We can represent a graph with a weighted adjacency matrix, where matrix value *W_i,j_* is the weight of the edge leading from node *i* to node *j* (absent = “-”). The weighted adjacency matrix of *Graph 1* is:

|  | P_1_ | M_F_ | P_2_ | M_S_ | P_3_ | M_B_ |
| --- | --- | --- | --- | --- | --- | --- |
| P_1_ | - | - | - | - | - | 15, 1 |
| M_F_ | 3, 1 | - | 3, 1 | - | - | - |
| P_2_ | - | - | - | - | - | 15, *n* |
| M_S_ | - | - | - | - | 12, 1 | - |
| P_3_ | - | - | - | - | - | 15, 1 |
| M_B_ | - | - | 15, *n* | - | - | - |

The minimum representation of this matrix is the smallest (i.e., most compressed) format that allows for an unambiguous reconstruction of the original graph structure. If all edge weights are possible at every edge, then the minimum unambiguous representation of the weighted adjacency matrix is the matrix itself.

However, for any particular dataset, we can survey the empirical diversity of graph structures and assume constant those features that do not vary across the dataset. We can use these assumptions to unambiguously decompress a simplified vector representation of a more complex graph structure. To illustrate this process (in reverse) for our dataset, consider a minimum representation of *Graph 1* as a vector of three character states.

| D | A | E |
| --- | --- | --- |

[1] Assume character states D, A, and E correspond to character state vectors (3, 1), (3, *n*), and (12, 1), respectively. These states are vectors of two elements. Note, though, that the character state vectors are different than the edge weight vectors in *Graph 1*. Like the first value of an edge weight vector, the first value of a character state vector represents the first time a plumage patch appears, in months. However, unlike the second value of an edge weight vector, the second value of a character state vector represents the *total* number of times a plumage patch is re/generated. Therefore, the second-index value for character states must equal the *sum of the second-index values of edge weights* leading to a given plumage node. This information will matter later.

| 3, 1 | 3, *n* | 12, 1 |
| --- | --- | --- |

[2] Assume character states at indices 1, 2, and 3 correspond to characters P_1_, P_2_, and P_3_, respectively.

| P_1_ | P_2_ | P_3_ |
| --- | --- | --- |
| 3, 1 | 3, *n* | 12, 1 |

[3] Assume there are exactly three molt nodes (M_F_, M_S_, and M_B_).

|  | P_1_ | M_F_ | P_2_ | M_S_ | P_3_ | M_B_ |
| --- | --- | --- | --- | --- | --- | --- |
| P_1_ | ? | ? | ? | ? | ? | ? |
| M_F_ | ? | ? | ? | ? | ? | ? |
| P_2_ | ? | ? | ? | ? | ? | ? |
| M_S_ | ? | ? | ? | ? | ? | ? |
| P_3_ | ? | ? | ? | ? | ? | ? |
| M_B_ | ? | ? | ? | ? | ? | ? |

[4] Assume the only edges present in the graph connect molt nodes and plumage nodes. In other words, there are no molt-molt edges, and no plumage-plumage edges.

|  | P_1_ | M_F_ | P_2_ | M_S_ | P_3_ | M_B_ |
| --- | --- | --- | --- | --- | --- | --- |
| P_1_ | - | ? | - | ? | - | ? |
| M_F_ | ? | - | ? | - | ? | - |
| P_2_ | - | ? | - | ? | - | ? |
| M_S_ | ? | - | ? | - | ? | - |
| P_3_ | - | ? | - | ? | - | ? |
| M_B_ | ? | - | ? | - | ? | - |

[5] Assume M_F_ exclusively generates plumage patches at 3 months, and M_S_ exclusively generates plumage patches at 12 months. All other plumage patches are generated via M_B_, which only produces plumages at annual intervals offset by 3 months (e.g., 15, 27, 39 months). Thus, a plumage patch that first appears at 3 months must have a (3, 1) edge leading from M_F_, and so on.

|  | P_1_ | M_F_ | P_2_ | M_S_ | P_3_ | M_B_ |
| --- | --- | --- | --- | --- | --- | --- |
| P_1_ | - | ? | - | ? | - | ? |
| M_F_ | 3, 1 | - | 3, 1 | - | - | - |
| P_2_ | - | ? | - | ? | - | ? |
| M_S_ | - | - | - | - | 12, 1 | - |
| P_3_ | - | ? | - | ? | - | ? |
| M_B_ | ? | - | ? | - | ? | - |

[6] Assume preformative (M_F_) and presupplemental molts (M_S_) only generate patches, never removing them. Thus, there are no edges leading to either of those nodes.

|  | P_1_ | M_F_ | P_2_ | M_S_ | P_3_ | M_B_ |
| --- | --- | --- | --- | --- | --- | --- |
| P_1_ | - | - | - | - | - | ? |
| M_F_ | 3, 1 | - | 3, 1 | - | - | - |
| P_2_ | - | - | - | - | - | ? |
| M_S_ | - | - | - | - | 12, 1 | - |
| P_3_ | - | - | - | - | - | ? |
| M_B_ | ? | - | ? | - | ? | - |

[7] Assume the prebasic molt (M_B_) is complete (i.e., removes all existing plumage patches before generating any new ones). Therefore, a plumage patch with appearance time *t* must have an edge leading to M_B,_ where the first-index value of that edge weight is equal to the lowest value from the cycle of prebasic molts (15, 27, 39…) that is also greater than *t*.

|  | P_1_ | M_F_ | P_2_ | M_S_ | P_3_ | M_B_ |
| --- | --- | --- | --- | --- | --- | --- |
| P_1_ | - | - | - | - | - | 15, 1 |
| M_F_ | 3, 1 | - | 3, 1 | - | - | - |
| P_2_ | - | - | - | - | - | 15, *n* |
| M_S_ | - | - | - | - | 12, 1 | - |
| P_3_ | - | - | - | - | - | 15, 1 |
| M_B_ | ? | - | ? | - | ? | - |

[8] Assume there are no plumage patches which reappear after an absence. Recalling [1], we know the second-index value of the character state vectors must equal the sum of the second-index values of the edge weight vectors leading to a given plumage nodes. We can now fill in the final edges leading from M_B_.

|  | P_1_ | M_F_ | P_2_ | M_S_ | P_3_ | M_B_ |
| --- | --- | --- | --- | --- | --- | --- |
| P_1_ | - | - | - | - | - | 15, 1 |
| M_F_ | 3, 1 | - | 3, 1 | - | - | - |
| P_2_ | - | - | - | - | - | 15, *n* |
| M_S_ | - | - | - | - | 12, 1 | - |
| P_3_ | - | - | - | - | - | 15, 1 |
| M_B_ | - | - | 15, *n* | - | - | - |

Assumptions [1] through [8] thus allow us to unambiguously reconstruct the weighted adjacency matrix (and therefore the complete structure) of *Graph 1*, given only the compressed vector of three character states. With these same assumptions, the characters and character states in our full analysis allowed us to unambiguously represent the plumage maturation processes for extant and ancestral manakin lineages. Note that some of the numbered assumptions simply refer to basic biology. For example, assumption [4] follows from biological definitions of “plumage” and “molt.” Other assumptions refer to the empirical diversity of graph structures in our specific dataset. For example, assumption [3] would *not* apply if our dataset featured even one taxon with a prealternate molt (M_A_), which occurs at annual intervals offset from M_B_ by ~6 months (Wolfe et al. 2010). No manakins have prealternate molts (Johnson and Wolfe 2018). We can thus apply assumption [3], which, in combination with assumptions [1] and [8], allows us to fill in the edges leading from M_B_ in the final step.

Viewing this process optimistically, one can define character states with respect to the empirical diversity of a clade in a way that represents changes to a complex underlying phenomenon (e.g., a graph structure) in a highly compressed format (e.g., a simplified state vector). More pessimistically, the definition of character states constrains the kinds of evolution that are possible to represent in a given model. In our study, we use assumptions about graph structures that make it impossible to represent the evolution of a prealternate molt. In an event-pairing analysis, evolution is assumed to involve only changes to the edges relating fixed nodes (Bininda-Emonds et al. 2002). In continuous comparative analyses of e.g., body size, there is only a single homologous character with a shifting state. One thus assumes a single, fixed node with an evolving node state, or a single, fixed edge with an evolving edge state, or, perhaps more generously, an entire graph measured only in terms of some holistic feature (e.g., the sum of all edge weights in a graph with an agnostic structure).

A general, formal method for phylogenetic comparative analyses of graph structures does not yet exist. Such a method would empower studies of biological graphs such as gene trees (Maddison 1997), cell type trees (Mah and Dunn 2024), synteny networks (Ghiurcuta and Moret 2014), gene regulatory networks (Levine and Davidson 2005), cell signaling networks (Stadtmauer et al. 2024), and anatomical body plans (Tarasov 2019), not to mention social networks, food webs, and so on.

Many practical questions remain. For example, which models best describe the evolutionary dependencies between edges in a particular set of graphs? In our analysis, we assume the dynamics of evolution are completely independent in different parts of the plumage-molt graph. Therefore, we allow separate transitions (and even separate transition models) for each character. In contrast, phylogenetic analyses of nucleic acid sequences—conceivably, a linear graph with shifting nodes but a fixed edge structure that defines locus position—assume that substitutions in one part of the graph inform substitutions in other parts, either universally (e.g., JC, GTR models; Arenas 2015), or in a small number of partitions (e.g., +G models; Yang 1994).

**Manakin plumage maturation descriptions**

Here, we provide brief accounts of male plumage maturation for 40 manakin taxa. These descriptions were translated into sets of characters (i.e., plumage patches) and character states (i.e., developmental schedules) for phylogenetic analysis. Each account is divided into three sections:

1. **Summary of male plumage maturation**. Plumage and molt terminology follows modified Wolfe-Ryder-Pyle conventions (Wolfe et al. 2010; Johnson and Wolfe 2018), with “definitive” used to specify the indefinitely repeated plumage cycle (Howell and Pyle 2015). Plumage and molt abbreviations: FCJ = first cycle juvenile plumage, FPF = first cycle preformative molt; FCF = first cycle formative plumage; SPB = second cycle prebasic molt; SCB = second cycle basic plumage; TPB = third cycle prebasic molt; TCB = third cycle basic plumage; DPB = definitive cycle prebasic molt; DCB = definitive cycle basic plumage. Here, we use codes to directly describe plumages and molts (e.g., “FPF is partial” = “the first cycle preformative molt is partial”).
2. **References and justification for plumage sequence description**. Data quality differed between taxa. We assigned three data quality categories: [A] Detailed maturation descriptions from published records, including banding studies with recaptured individuals (n = 20 taxa). [B] Limited or partially conflicting information from published records, including banding studies with recaptured individuals, supplemented by unpublished banding records or additional museum specimens for this study (n = 15 taxa). [C] Limited descriptions from museum specimens only (n = 5 taxa). Museum abbreviations: AMNH = American Museum of Natural History; YPM = Yale Peabody Museum of Natural History.
3. **Photo examples**. Photograph vouchers from the Macaulay Library (ML) at the Cornell Lab of Ornithology. Note that photos cannot be assigned to specific plumage or molts (e.g., if male FCF is identical to female DCB, we can only present a photo example for “female” plumage).

We make several key assumptions in mapping plumage maturation across species (Schaedler et al. 2021). In particular, we assume that FPF is partial for all species, that basic molts are complete for all species, and that molt timing is consistent across species (FPF ~3 months, with subsequent basic molts at ~12-month intervals). These features are generalized from detailed records for a subset of species in Johnson and Wolfe (2018) and Scholer et al. (2021). We note where additional studies could provide better information about plumage maturation in some accounts below.

Our goal is to establish a proper dataset for comparative analysis by compiling and supplementing existing data on male plumage maturation in manakins. We do not aim to provide complete descriptions of definitive plumages. For full plumage descriptions, readers can consult Kirwan and Green (2011) or species accounts in Cornell’s Birds of the World. Finally, we do not characterize the intraspecific variation reported for many taxa (Ryder and Durães 2005; Johnson and Wolfe 2018; Schaedler et al. 2021; Scholer et al. 2021). Future studies of this intraspecific variation in manakins would greatly aid our understanding of delayed plumage maturation and its evolution.

*Tyranneutes virescens*

Sexually monochromatic. (FCF) female plumage with retained juvenile wing feathers. (DCB) female plumage.

Data quality [B]. Ribeiro et al. (2015) describe as “nearly monochromatic.” Schaedler et al. (2021) describe this taxon as sexually dichromatic with delayed maturation of male definitive yellow crown until DCB. This description follows Johnson and Wolfe (2018), who suggest subtle, quantitative sexual differences in the extent of the yellow crown and describe FCF as having yellow crown “partially concealed or absent” (i.e., presumably reduced with respect to DCB). In AMNH collection (~6 female-tagged, ~25 male-tagged), all except one female-tagged specimen have some yellow on the crown, and all male-tagged specimens have yellow on the crown. There is some variation in prominence, with female-tagged specimens appearing to have less prominent yellow crowns. These specimens suggest variation not in terms of the area of the yellow crown, but rather in the extent of the greenish feather tips within the crown. Given the lack of clear sexual dichromatism among definitive plumages, and the additionally limited evidence of predefinitive variation, we revise the description from Schaedler et al. (2021) to classify this taxon as sexually monochromatic with no delayed plumage maturation. Additional studies are needed to quantify sex- and age-related patterns in the yellow crown.

Definitive: ML194703421

*Neopelma aurifrons*

Sexually monochromatic. (FCF) female plumage with retained juvenile wing feathers. (DCB) female plumage.

Data quality [B]. Schaedler et al. (2021) describe as sexually monochromatic with no delayed plumage maturation, following description in Kirwan and Green (2011). Additional studies are needed to fully describe the timing and extent of FPF.

Definitive: ML262033451

*Neopelma pallescens*

Sexually monochromatic. (FCF) female plumage with retained juvenile wing feathers. (DCB) female plumage.

Data quality [B]. Ribeiro et al. (2015) describe as monochromatic. Schaedler et al. (2021) follow description in Kirwan and Green (2011). Ferreira and Lopes (2018) similarly describe FCJ of recent fledglings as overall similar to sexually monochromatic DCB.

Definitive: ML205104151, ML205323361

*Neopelma chrysocephalum*

Sexually monochromatic. (FCF) female plumage with retained juvenile wing feathers. (DCB) female plumage.

Data quality [B]. Ribeiro et al. (2015) describe as monochromatic. Johnson and Wolfe (2018) describe no clear sexual dichromatism and no delayed plumage maturation (although note data provided from one individual). Additional studies are needed to quantify sex- and age-related patterns in the extent of the concealed yellow crown. AMNH collections suggest all individuals have warm yellow crowns, with no clear differences between female-tagged and male-tagged specimens.

Definitive: ML405077991, ML404620241

*Masius chrysopterus*

Sexually dichromatic. (FCF) female plumage with retained juvenile wing feathers. (SCB) plus female wing feathers, yellow crown accentuated at the forecrown, flattened and darkened nape, and appearance of some black body feathers. (DCB) male definitive, including black body with yellow crown, yellow throat, yellow patches hidden in folded wings and tail, and scaley, tobacco-colored nape.

Data quality [B]. Schaedler et al. (2021) follow Taylor et al. (2020) in describing three predefinitive plumage stages with uncertain timing. Independent banding records from D. McDonald and N. Büttner (*pers. comm.*) for separate sites in Ecuador support the annual timing of each stage and partial FPF, although formal description of delayed plumage maturation in this species is still outstanding. The asymmetrical yellow edging that forms hidden patches in both the wings and tail in this taxon (e.g., ML391279521) has a distinct identity compared to the restricted, symmetrical white wing patch in *Corapipo*, and we code it as a separate character.

Female: ML449827661, ML133660421

Male predefinitive: ML608667835, ML204890431

Male definitive: ML588959091, ML588959111, ML521155321

*Corapipo gutturalis*

Sexually dichromatic. (FCF) female plumage with retained juvenile wing feathers. (SCB) plus female wing feathers, white throat, and black mask. (DCB) male definitive, with black body, white throat, and hidden white wing patch.

Data quality [A]. Schaedler et al. (2021) follow descriptions in Prum (1986), Johnson and Wolfe (2018), and Aramuni (2019). Johnson and Wolfe (2018) provide detailed records showing the delay of the white wing patch until DCB.

Female: ML204133411, ML368797241

Male predefinitive: ML363146471, ML473853741, ML508906011

Male definitive: ML204133311

*Corapipo altera*

Sexually dichromatic. (FCF) female plumage with retained juvenile wing feathers. (SCB) plus female wing feathers, white throat, and black mask. (DCB) male definitive, with black body and white throat.

Data quality [A]. Schaedler et al. (2021) follow description in Jones et al. (2014), which aligns with descriptions for *C. gutturalis*.

Female: ML174527761

Male predefinitive: ML363351401, ML381062901

Male definitive: ML143970211

*Corapipo leucorrhoa*

Sexually dichromatic. (FCF) female plumage with retained juvenile wing feathers. (SCB) plus female wing feathers, white throat, and black mask. (DCB) male definitive, with black body and white throat.

Data quality [B]. Schaedler et al. (2021) follow description in Rosselli (1994), which aligns with descriptions for *C. gutturalis*.

Female: ML198145241, ML80524531

Male predefinitive: ML540613271, ML191658641, ML531466031

Male definitive: ML543551161

*Chiroxiphia caudata*

Sexually dichromatic. (FCF) female plumage with retained juvenile wing feathers, plus restricted red crown. (FCS) plus black mask. (SCB) plus female wing feathers, some blue body feathers, and black head. (DCB) male definitive, with red crown, black head, blue body, and black wings and tail.

Data quality [A]. Schaedler et al. (2021) follow description from Mallet-Rodrigues and Dutra (2012). Additional studies are needed to identify the exact timing of the black mask, although Mallet-Rodrigues and Dutra (2012) make clear that both the red cap and black face feathers appear in the first cycle (i.e., either FCF or FCS). Here, we assume the mask is generated via FPS as described for *C. lanceolata* (DuVal 2005). As Schaedler et al. (2021) discuss, closer examination of this species may shed light on the extent to which additional plumage stages, such as those described in Mallet-Rodrigues and Dutra (2012), may be staggered between molts.

Female: ML612262709

Male predefinitive: ML603127111, ML589071901, ML474193321, ML92416921

Male definitive: ML269492221

*Chiroxiphia boliviana*

Sexually dichromatic. (FCF) female plumage with retained juvenile wing feathers, plus restricted red crown. (FCS) plus black mask. (SCB) plus female wing feathers, blue mantle, and additional patches of black body. (DCB) male definitive, with red crown, blue mantle, and black body.

Data quality [A]. For this study, we assign plumage maturation in *C. pareola* as fully identical to *C. lanceolata*. Scholer et al. (2021) follow Kirwan and Green (2011) in simply asserting the presence of delayed plumage maturation with uncertain timing and aspect for predefinitive plumage stages. Scholer et al. (2021) provide additional details by clarifying FCF includes a red crown, uncertain and variable amounts of black across the body, and (less well described for first cycle plumages in the genus) sometimes blue feathers on the back, SCB with more substantial blue and black feathers, and third cycle molt (=DPB) that results in the male definitive plumage. M. Villegas and B. Loiselle (*pers. comm*.) support these stages with additional field records, suggesting that plumage maturation in this taxon parallels *C. lanceolata*, including black feathers appearing within the first cycle, in contrast to the longer maturation in *C. linearis*. Here, we assume the mask is generated via FPS as described for *C. lanceolata* (DuVal 2005). Additional studies are needed to investigate the possibility of blue mantle feathers appearing in FCF (as also hinted at for *C. pareola*, but not described in *C. lanceolata* or *C. linearis*), as well as characteristics of FPS.

Female: ML37257081

Male predefinitive: ML274857091, ML72636041, ML58131691

Male definitive: ML612929239, ML62914531

*Antilophia galeata*

Sexually dichromatic. (FCF) female plumage with retained juvenile wing feathers. (SCB) plus tufted forecrown with individual red feathers on cap and onto mantle, concentrated black mask, concentrated black on the proximate ends of tail feathers and upper wings, and streaky black across the back, belly, and throat. (DCB) male definitive, with black body and red crown extending into cape.

Data quality [B]. Schaedler et al. (2021) follow Allen (1893), Marini and Cavalcanti (1992), and Kirwan and Green (2011) in describing some young males with red along the crown and mantle and/or scattered black across the body. AMNH collections support two predefinitive plumage classes for male-tagged specimens: (A) fully female plumages, sometimes with a small number of individual, bright red feathers (AMNH58533, AMNH33481, AMNH33514, AMNH33512, AMNH33561, AMNH33485, AMNH492569, AMNH33493) and (B) more extensive red crown and variable amounts of black feathers, with black mostly concentrated around face, base of tail, upper wings, and patchier across throat and rest of body (AMNH58514, AMNH33488, AMNH33516, AMNH33494, AMNH33632, AMNH33556, AMNH33557, AMNH33499, AMNH33559, AMNH33476, AMNH33554, AMNH33549). At least one male-tagged specimen is intermediate between these stages (AMNH33485), and no specimens featured clear information on molt or skull ossification. Close examination of male-tagged specimens in the first, more female stage revealed a subtle, shimmering, chestnut-red aspect to the mantle that may be difficult to observe in the field and is unlikely to function as a biological signal. L. N. Kajiki (*pers. comm.)* generally supports the relative timing and aspect of these two predefinitive plumage stages from banding records of ~250 individuals, but formal analysis of these records—and a more detailed description of FPF and plumage maturation more generally—is desperately needed.

Female: ML552539961

Male predefinitive: ML204745391, ML205536581, ML81436311, ML267009441, ML509584571, ML421373261, ML244323021, ML60314451, ML615257825, ML245376591

Male definitive: ML109497121

*Chiroxiphia pareola*

Sexually dichromatic. (FCF) female plumage with retained juvenile wing feathers, plus restricted red crown. (FCS) plus black mask. (SCB) plus female wing feathers, blue mantle, and additional patches of black body. (DCB) male definitive, with red crown, blue mantle, and black body.

Data quality [A]. Schaedler et al. (2021) follow description in Cárdenas‐Posada et al. (2018). Subspecies *regina* has yellow, not red, crown. Additional studies are needed to identify the exact timing of the black mask, although Cárdenas‐Posada et al. (2018) make clear that both the red cap and black face feathers appear “during the second year” (i.e., first molt cycle, either FCF or FCS) rather than the second molt cycle as in *C. linearis*. J. Blake (*pers. comm.*) supports this description, but information on exact molt timing remains limited and additional studies are needed to identify differences between *C. pareola* and *C. lanceolata*. Here, we assume the mask is generated via FPS as described for *C. lanceolata* (DuVal 2005). Additional studies should also clarify the extent to which young males gain a blue aspect on the mantle prior to substantial black feathers, unlike both *C. lanceolata* and *C. linearis* (e.g., ML389739791).

Female: ML343460301, ML245630611,

Male predefinitive: ML417429301, ML194035601, [*regina*] ML111236681, ML611838422, ML282213291

Male definitive: ML205094531, ML77577211

*Chiroxiphia linearis*

Sexually dichromatic. (FCF) female plumage with retained juvenile wing feathers, plus restricted red crown. (SCB) plus female wing feathers and black mask. (TCB) plus blue mantle and more substantial black body feathers. (DCB) male definitive, with red crown, blue mantle, and black body.

Data quality [A]. Schaedler et al. (2021) follow detailed description from Doucet et al. (2007). Note the clear description of the timing of molt, including the three-year delay before male definitive plumage and the “black face” stage via SPB (rather than FPS as in *C. lanceolata*).

Female: ML290665661, ML200117761

Male predefinitive: ML613865439, ML152290181, ML71359721, ML100076281, ML226077641, ML156641571

Male definitive: ML563499741, ML563483951

*Chiroxiphia lanceolata*

Sexually dichromatic. (FCF) female plumage with retained juvenile wing feathers, plus restricted red crown. (FCS) plus black mask. (SCB) plus female wing feathers, blue mantle, and additional patches of black body. (DCB) male definitive, with red crown, blue mantle, and black body.

Data quality [A]. Schaedler et al. (2021) follow detailed description from DuVal (2005). Note the clear description of the timing of molt, including the apparent first-cycle supplemental molt in the “black face” stage ~12 months after hatching (DuVal 2005).

Female: ML189427131, ML173383301

Male predefinitive: ML204927821, ML327293981, ML336393221, ML489081781, ML532585461

Male definitive: ML139196261, ML545266801

*Xenopipo uniformis*

Sexually monochromatic. (FCF) female plumage with retained juvenile wing feathers. (DCB) female plumage.

Data quality [B]. Ribeiro et al. (2015) describe as monochromatic. Schaedler et al. (2021) follow description in Snow (2020a) to assert no delayed plumage maturation. Additional studies are needed to clarify the extent of FPF.

Definitive: ML205203411, ML173298961

*Xenopipo atronitens*

Sexually dichromatic. (FCF) female plumage with retained juvenile wing feathers. (DCB) male definitive, with black body.

Data quality [C]. Schaedler et al. (2021) follow descriptions in Kirwan and Green (2011) to loosely assert the presence of delayed plumage maturation with young males described in variable transitional plumages (actively molting?) with small amounts of black. Museum collections support the presence of a male predefinitive stage given male-tagged specimens in either an entirely female, green plumage (AMNH461489, AMNH274644, YPM101657 testes 4x5 mm skull 100% ossified) or with scant, restricted black patches on an otherwise female plumage (AMNH311127, AMNH311111, AMNH311112, AMNH274893, AMNH311114, AMNH311116, AMNH311113). Additional studies are needed to confirm the extent of FPF and clarify the variation and timing of patchy black elements in young male plumages (e.g., ML588896331).

Female: ML204137161

Male definitive: ML530765741

*Chloropipo unicolor*

Sexually dichromatic. (FCF) female plumage with retained juvenile flight feathers. (DCB) male definitive, with black body.

Data quality [B]. Schaedler et al. (2021) follow Kirwan and Green (2011) to assert delayed plumage maturation, with descriptions from the latter suggesting some young male plumages are generally female but can feature scattered black feathers. AMNH collections provide further clarification, with male-tagged specimens in fully female plumage (AMNH820305: testes 1 mm; AMNH820128 testes 2 mm, 1 mm; AMNH820679, testes 6x3 mm, 4x3 mm). These specimens still provide uncertain data on male FCF, because the skulls of all three are tagged as unossified or incompletely ossified. Based on skull development in other manakins (Johnson and Wolfe 2018), ossified skulls would be needed to confirm FCF rather than FCJ (i.e., if ossification occurs during FCF, no FCJ manakins should have ossified skulls, although some FCF manakins may have unossified skulls). However, the same AMNH collection (1968–1970, J. Weske) has puzzling skull notes, including two male-tagged specimens in full male DCB that are tagged with incomplete skulls (AMNH820693, AMNH820681). Additional studies are needed to identify the timing and extent of FPF and the aspect of male FCF, including sex- and age-related differences in white underwing coverts (e.g., green-plumaged birds with white underwings: ML546919191).

Female: ML546404211

Male definitive: ML557834441

*Chloropipo flavicapilla*

Minor sexual dichromatism. (FCF) female plumage, included greener or dully yellow head, with retained juvenile wing feathers. (DCB) male definitive, with yellower head especially concentrated on crown.

Data quality [B]. Ribeiro et al. (2015) describe as “nearly monochromatic.” Schaedler et al. (2021) follow descriptions in Hellmayr (1929) and Kirwan and Green (2011) to assert male FCF has reduced contrast in the yellow head akin to female DCB. Additional studies are needed to quantify sex- and age-related patterns in the extent of the yellow head, along with timing and extent of FPF.

Female: ML205026451, ML205016461

Male definitive: ML205129261, ML616273315

*Cryptopipo holochlora*

Sexually monochromatic. (FCF) female plumage with retained juvenile wing feathers. (DCB) female plumage.

Data quality [A]. Scholer et al. (2021) provide detailed information on *Cryptopipo holochlora viridor* from southeast Peru. They report retained juvenile flight feathers and only slight differences in color between FCF and DCB, the former being slightly duller with additional wear on the retained juvenile feathers.

Definitive: ML484089091

*Cryptopipo litae*

Sexually monochromatic. (FCF) female plumage with retained juvenile wing feathers. (DCB) female plumage.

Data quality [B]. Ribeiro et al. (2015) describe *Xenopipo holochlora* (=*Cryptopipo holochlora* unk. subspp., prior to split between *C. holochlora* and *C. litae*). Schaedler (2021) follows Wetmore (1972) in stating *Cryptopipo holochlora* has no delayed plumage maturation, based on the description of no differences in “immature” (FCJ?) birds other than “somewhat duller colored.” The description in Wetmore (1972) applies to *Chloropipo holochlora litae* (=*Cryptopipo litae litae*) and *Chloropipo holochlora suffusa* (=*Cryptopipo litae suffusa*). Additional studies are needed to clarify the timing and extent of FPF in this taxa, as opposed to *C. holochlora*.

Definitive: ML120022311

*Lepidothrix serena*

Sexually dichromatic. (FCF) female plumage with retained juvenile wing feathers. (DCB) male definitive, with black body, white forecrown, blue rump, and rich yellow belly and breast spot.

Data quality [A]. Schaedler et al. (2021) offer an uncertain description based on Prum (1994) and Snow (2020b), missing the clearer description from Johnson and Wolfe (2018). Although the female definitive plumage also features a yellow belly and partially yellow breast, we code the richer, high-contrast yellow belly and breast spot in male DCB as distinct male patches.

Female: ML204139181

Male definitive: ML204135401

*Lepidothrix coeruleocapilla*

Sexually dichromatic. (FCF) female plumage with retained juvenile wing feathers. (DCB) male definitive, with black body, blue crown, and blue rump.

Data quality [A]. Schaedler et al. (2021) offer an uncertain description following Kirwan and Green (2011). Scholer et al. (2021) describe both FCF and DCB from banding data.

Female: ML130641861, ML419332031

Male definitive: ML419654791, ML375850771

*Lepidothrix velutina*

Sexually dichromatic. (FCF) female plumage with retained juvenile wing feathers. (DCB) male definitive, with black body and blue crown.

Data quality [C]. Limited information from a small number of museum specimens and comparisons to closely related species. Snow and Kirwan (2020) apply the plumage and molt description from the split *L. coronata* as described by Ryder and Durães (2005). AMNH collections include one male-tagged specimen in a fully female plumage (AMNH390870) and another male-tagged specimen in a female plumage with the addition of a few bright blue feathers running in a short, broken line from above right eye to the forecrown (AMNH272086). Consistent with other taxa formerly assigned to the *L. coronata* species complex, these specimens support the presence of delayed plumage maturation with a simple female predefinitive plumage in the first cycle, although some details are difficult to track given various related taxa with green-bodied (rather than black-bodied) males (Kirwan and Green 2011). Additional studies are needed to understand the specifics of FPF and male FCF.

Female: ML615825929

Male definitive: ML142752011

*Lepidothrix coronata coronata*

Sexually dichromatic. (FCF) female plumage with retained juvenile wing feathers. (DCB) male definitive, with black body and blue crown.

Data quality [B]. Schaedler et al. (2021) describe *L. coronata* as having two predefinitive plumage stages—female FCF followed by SCB with variable amounts of blue in the crown and black on the body. This description is based on two kinds of uncertain reports: (1) Ryder and Durães (2005), who somewhat vaguely report males retaining some level of female plumage into the second cycle, and (2) Kirwan and Green (2011), who draw on the preceding source and additionally describe some museum specimens with male crown feathers on a green body. Schaedler et al. (2021) interpret these descriptions as providing evidence for a distinct male SCB with a green/black body and a blue crown. This interpretation potentially confounds (A) variation of FCF, (B) poorly detailed intraspecific variation in the time and extent of SPB and/or the aspect of SCB, and (C) limited information on sub/species diversity within the *L. coronata* species complex, which features some taxa with green, rather than black, bodies in male DCB (Moncrieff et al. 2022). Counter to the description in Schaedler et al. (2021), there appears to be no evidence of predefinitive males with a green body plus blue crown for *L. c. coronata*, a taxon in which male DCB have a black body. AMNH collections feature three male-tagged specimens with female plumages (AMNH256400, AMNH239450, AMNH816762 testes 2x1 mm). Two of these specimens (AMNH239450, AMNH816762) have scant individual blue crown feathers and (on close inspection) a shimmery, blue-green wash to the crown, hindcrown, and/or back, but no actual crown. Based on the lack of clear description for multiple distinct predefinitive plumage stages, and the fact that the present study does not incorporate intraspecific variation (as importantly described by Ryder and Durães 2005), we conservatively classify this species as having a single, female predefinitive plumage stage. Additional studies are needed to clarify taxon-, population-, and even intrapopulation-level variation in male predefinitive plumages, with explicit attention towards corresponding variation in male DCB.

Female: ML158318931 [Ecuador]

Male definitive: ML208351561 [Ecuador]

*Lepidothrix coronata caelestipileata*

Sexually dichromatic. (FCF) female plumage with retained juvenile wing feathers. (DCB) male definitive, with green (i.e., female) body and wings but with a black head and blue crown.

Data quality [A]. Scholer et al. (2021) describe both FCF and DCB from banding data. As discussed in the above description for *L. c. coronata*, additional studies are needed to understand FCF variation between green- and black-bodied *L. coronata* lineages.

Female: ML610586553 [Madre de Dios, Peru], ML609828509 [uncertain taxon—La Paz, Bolivia]

Male definitive: ML380224341, ML205973951 [both Madre de Dios, Peru]

*Heterocercus flavivertex*

Sexually dichromatic. (FCF) female plumage with retained juvenile wing feathers. (DCB) male definitive, overall similar to female plumage but with rich brown underparts, high-contrast white throat, and hidden yellow crown.

Data quality [B]. Schaedler et al. (2021) follow description in Prum et al. (1996), plus consistent details in Kirwan and Green (2011). AMNH collections feature four male-tagged specimens in female plumages (AMNH276300, AMNH177636, AMNH493643, AMNH433413).

Female: ML581570391, ML183512641

Male definitive: ML204137181, ML453453521

*Manacus manacus*

Sexually dichromatic. (FCF) female plumage with retained juvenile wing feathers. (DCB) male definitive, with black cap, black back, gray belly, gray rump, and white collar wrapping around neck.

Data quality [A]. Schaedler et al. (2021) follow description in Johnson and Wolfe (2018), with corresponding details in Tu et al. (2020). Morales-Betancourt and Castaño-Villa (2018) description quantitative UV reflectance differences in male predefinitive plumages, but these do not suggest qualitative sexual differences in plumage patches and are not included in the present study. Additional studies are needed to clarify the details and biological relevance of UV reflectance differences in these and other manakins.

Female: ML451089331, ML205389311

Male definitive: ML448955081, ML56898711

*Manacus candei*

Sexually dichromatic. (FCF) female plumage with retained juvenile wing feathers. (DCB) male definitive, with black cap, black flight feathers, white collar wrapping around neck, and rich yellow belly, although continuing with a female (albeit slightly richer yellow) back and rump.

Data quality [A]. Schaedler et al. (2021) follow description in Wolfe et al. (2009). Although the female plumage also features a yellow belly, we code the richer, high-contrast yellow belly in the male DCB as a distinct patch.

Female: ML418711511

Male definitive: ML549365041, ML613986469

*Manacus vitellinus*

Sexually dichromatic. (FCF) female plumage with retained juvenile wing feathers. (DCB) male definitive, with black cap, black flight feathers, and yellow collar wrapping around neck, although continuing with a female belly and rump.

Data quality [A]. Schaedler et al. (2021) follow description in Day et al. (2006).

Female: ML139195751, ML514867391

Male definitive: ML266148701, ML615825131

*Pipra filicauda*

Sexually dichromatic. (FCF) female plumage with retained juvenile wing feathers. (SCB) plus variable amounts of red on the cap, yellow in the face and underparts, and black on back including across wings and tail. (DCB) male definitive, with solid red cap extending into nape, yellow face and underparts, black back, and hidden white wing patch.

Data quality [B]. Schaedler et al. (2021) follow description in Ryder and Durães (2005), with consistent details in Kirwan and Green (2011), although Schaedler et al. (2021) neglect the variable black that can appear concurrent to carotenoid plumage elements in SCB. T. B. Ryder (*pers. comm.*) verifies that the hidden white wing patch appears via DPB, not SPB, and further emphasizes the substantial variation in extent of SCB, which can include carotenoid and black elements across the entire body (albeit still littered with green feathers).

Female: ML612770739

Male predefinitive: ML610118840

Male definitive: ML20783911, ML320133401

*Pipra fasciicauda*

Sexually dichromatic. (FCF) female plumage with retained juvenile wing feathers. (SCB) plus variable amounts of red wash on the head down to breast, yellow underparts, and black on back including across wings and tail. (DCB) male definitive, with red head washed down into yellow underparts, black back, and hidden white wing patch.

Data quality [A]. Schaedler et al. (2021) offer an uncertain description based on limited details in Robbins (1985) and Kirwan and Green (2011). Scholer et al. (2021) provide substantially clearer information based on banding data, including a description of the partial FPF and the intermediate SCB featuring washed red head, yellow underparts, black on wings, tail, and back, and reduced or absent white wing patches.

Female: ML352466861, ML371301091

Male predefinitive: ML113485871, ML554623261

Male definitive: ML384285991, ML352466831

*Pseudopipra pipra*

Sexually dichromatic. (FCF) female plumage with retained juvenile wing feathers. (SCB) plus gray head. (DCB) male definitive, with black body and white crown.

Data quality [A]. Schaedler et al. (2021) follow detailed description in Johnson and Wolfe (2018), drawn directly from thousands of banding records near Manaus, Brazil. Details about the relative extent of gray head in male FCF versus female DCB may be difficult to parse from verbal descriptions (e.g., Ryder and Durães 2005, discussing a different, potentially taxonomically uncertain lineage of *Pseudopipra*) or museum specimens (Kirwan and Green 2011; Berv et al. 2021). AMNH collections including females with various amounts of gray on the head, but also male-tagged specimens with more extensive gray. AMNH collections include no male-tagged specimens with a gray head and white crown as clearly described for some other *Pseudopipra* lineages. Additional studies are needed to clarify individual- or population-level variation in gray heads of male FCF versus female DCB.

Female: ML583319821 [Vaupés, Colombia], ML205188011 [French Guiana]

Male predefinitive: See SCB in Johnson and Wolfe (2018) account for *Dixiphia pipra pipra*

Male definitive: ML205807421 [Amapá, Brazil]

*Pseudopipra microlopha separabilis*

Sexually dimorphic. (FCF) female plumage with retained juvenile wing feathers. (SCB) plus a gray crown. (DCB) male definitive, with black body and white crown.

Data quality [C]. Schaedler et al. (2021) follow description in Berv et al. (2021), in turn informed by Zimmer (1936). Zimmer (1936) first noted the curious fact that predefinitive males have a gray, not white, crown. AMNH collections include male-tagged specimens in both fully female plumage (AMNH492921) and with the astounding gray crown (AMNH128721, AMNH128720). Additional studies are needed to verify the extent and timing of FPF, and to understand the evolutionary transition between gray and white crowns in this taxon versus e.g., *P. cephaleucos.*

Female: ML36368261 [Bahia, Brazil]

Male predefinitive: ML602227891 [Bahia, Brazil], ML217336361 [Bahia, Brazil]

Male definitive: ML36928921 [Bahia, Brazil]

*Pseudopipra cephaleucos*

Sexually dimorphic. (FCF) female plumage with retained juvenile wing feathers. (SCB) plus a white crown. (DCB) male definitive, with black body and white crown.

Data quality [C]. Schaedler et al. (2021) follow description in Berv et al. (2021), in turn informed by Zimmer (1936). Additional studies are needed to verify the extent and timing of FPF and the aspect of SCB, especially in contrast to local variation in gray-headed female DCB.

Female: ML568853051

Male predefinitive: Images not available

Male definitive: ML204874951

*Ceratopipra mentalis*

Sexually dimorphic. (FCF) female plumage with retained juvenile wing feathers. (DCB) male definitive, with red head, black body, and yellow leg tufts.

Data quality [A]. Schaedler et al. (2021) follow detailed description in Wolfe et al. (2009), with consistent details in Kirwan and Green (2011).

Female: ML379439271, ML535224561

Male definitive: ML613638644, ML434455941

*Ceratopipra chloromeros*

Sexually dimorphic. (FCF) female plumage with retained juvenile wing feathers. (DCB) male definitive, with red head, black body, and yellow leg tufts.

Data quality [A]. Schaedler et al. (2021) follow general description in Tello (2001). Scholer et al. (2021) provide detailed description from banding records.

Female: ML222410451

Male definitive: ML297064501

*Ceratopipra erythrocephala*

Sexually dimorphic. (FCF) female plumage with retained juvenile wing feathers. (DCB) male definitive, with yellow head, black body, and red and white leg tufts.

Data quality [A]. Schaedler et al. (2021) follow detailed description in Johnson and Wolfe (2018).

Female: ML205432191, ML51204711

Male definitive: ML207318021

*Machaeropterus deliciosus*

Sexually dichromatic. (FCF) female plumage with retained juvenile wing feathers. (DCB) male definitive, with red forecrown, chestnut brown body, black wings and tail, and visible white wing fringes.

Data quality [B]. Detailed banding records from Ecuador provided by D. Becker (*pers. comm.*) include multiple hatch year males first captured in a fully female plumage (or with only one or two male-like feathers) that are recaptured one year later (n = 3) or multiple years later (n = 1) in a fully male definitive plumage. Initial captures of all female birds were in December, whereas local breeding seasons peak between February and April, suggesting these female plumages were FCF and not FCJ (skull ossification not recorded). These records provide independent support for a female FCF in males, consistent with all other observations in *Machaeropterus*. Additional studies are needed to verify FPF and male FCF.

Female: ML206121961, ML307932011
Male definitive: ML114689001, ML557385541

*Machaeropterus pyrocephalus*

Sexually dichromatic. (FCF) female plumage with retained juvenile wing feathers. (DCB) male definitive, with yellow crown with central red stripe, chestnut brown body with streaks across the breast, and visible white wing patch.

Data quality [A]. Schaedler et al. (2021) assert the presence of delayed plumage maturation but offer an uncertain description, perhaps about molting birds, following Hilty (2003) and Kirwan and Green (2011). Scholer et al. (2021) offers a precise description from banding records.

Female: ML555050551, ML592259001

Male definitive: ML205150581, ML546507691

*Machaeropterus striolatus*

Sexually dichromatic. (FCF) female plumage with retained juvenile wing feathers. (DCB) male definitive, with red crown, and streaked, red-brown underparts.

Data quality [C]. Schaedler et al. (2021) follow Kirwan and Snow (2020) in asserting the presence of delayed plumage maturation. YPM collections feature five male-tagged specimens either in entirely female plumages (YPM183855, YPM183860, YPM116848), or with a few, scattered, bright red feathers (YPM179587, YPM116745). Additional studies are needed to verify FPF and male FCF.

Female: ML276514861, ML121610751

Male definitive: ML208189541, ML392197101

**Alternative phylogeny**

Results were congruent between the main phylogenetic analysis using the Harvey et al. (2020) tree and the alternative analysis using the Leite et al. (2021) tree (Fig. S1, Table S1). Given the smaller number of taxa used for the alternative analysis, there was additional uncertainty regarding low-confidence states for the ancestors of Pipridae (node 1 in Table S1), Piprinae (node 4), and Ilicurini (node 5). This analysis further supported a third independent origin of prolonged, two-year delayed plumage maturation in *Pseudopipra*, which is sister to *Ceratopipra* in the Leite tree (Fig. S1).

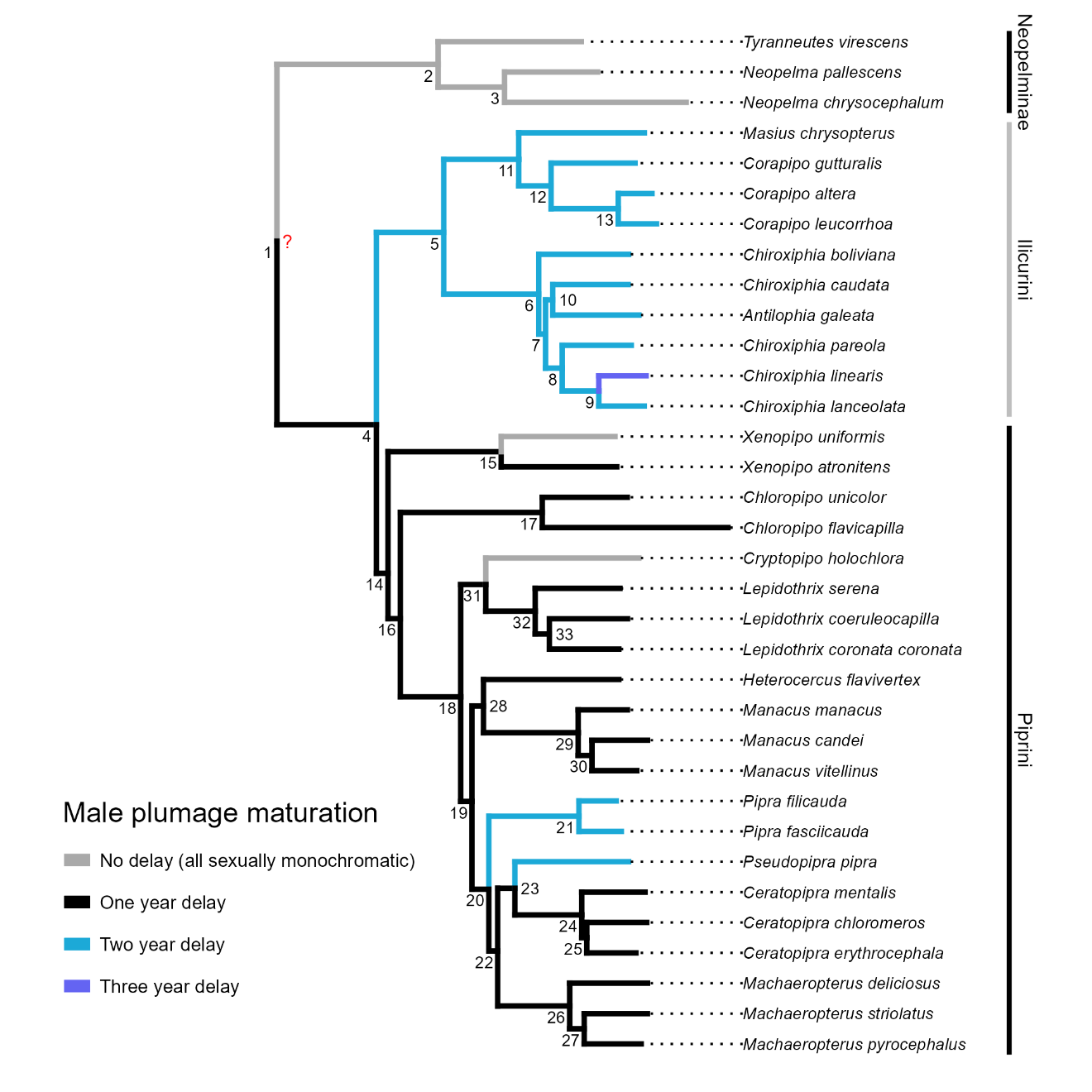

**Figure S1**. Alternative phylogenetic history of male plumage maturation in manakins. Tree from Leite et al. (2021). Edges colored based on the reconstruction of developmental schedules across a set of individual plumage patch characters at each child node. Red question mark indicates uncertainty surrounding plumage maturation at the root node. Node numbers correspond to Table S1.

**Table S1**. Ancestral male plumage maturation schedules given the Leite manakin phylogeny. Maximum likelihood character states shown for each of 28 plumage patch characters. Uncertain states indicated in gray (scaled likelihood <0.75) or censored as “?” (scaled likelihood <0.50). Absent characters (state ϕ) shown with a dash. Node numbers correspond to Figure S1.

|  |  |  |  |  |  |  | **Character** | | | | | | | | | | | | | | | | | | | | | | | | |
| --- | --- | --- | --- | --- | --- | --- | --- | --- | --- | --- | --- | --- | --- | --- | --- | --- | --- | --- | --- | --- | --- | --- | --- | --- | --- | --- | --- | --- | --- | --- | --- |
| **Node** |  | **Ancestor** |  | **1** | **2** | **3** | **4** | **5** | **6** | **7** | **8** | **9** | **10** | **11** | **12** | **13** | **14** | **15** | **16** | **17** | **18** | **19** | **20** | **21** | **22** | **23** | **24** | **25** | **26** | **27** | **28** |
| 1 |  | Pipridae (Root) |  | D | - | B | - | - | - | - | - | - | - | - | - | - | - | - | - | - | - | - | - | - | - | - | - | - | - | - | - |
| 2 |  | Neopelminae |  | A | B | - | - | - | - | - | - | - | - | - | - | - | - | - | - | - | - | - | - | - | - | - | - | - | - | - | - |
| 3 |  | *Neopelma* |  | A | B | - | - | - | - | - | - | - | - | - | - | - | - | - | - | - | - | - | - | - | - | - | - | - | - | - | - |
| 4 |  | Piprinae |  | D | - | B | - | - | - | - | - | - | - | - | - | - | - | - | - | - | - | - | - | - | - | - | - | - | - | - | - |
| 5 |  | Ilicurini |  | G | F | B | - | - | - | - | - | - | - | - | - | - | ? | - | - | - | - | - | - | - | - | - | - | - | - | - | - |
| 6 |  | *Chiroxiphia* |  | G | F | B | - | - | - | - | - | - | - | - | - | - | E | - | A | - | - | - | - | - | - | - | - | - | - | B | - |
| 7 |  |  |  | G | F | B | - | - | - | - | - | - | - | - | - | - | E | - | A | - | - | - | - | - | - | - | - | - | - | B | - |
| 8 |  |  |  | G | F | B | - | - | - | - | - | - | - | - | - | - | E | - | A | - | - | - | - | - | - | - | - | - | - | B | - |
| 9 |  |  |  | G | F | B | - | - | - | - | - | - | - | - | - | - | E | - | A | - | - | - | - | - | - | - | - | - | - | B | - |
| 10 |  |  |  | G | F | B | - | - | - | - | - | - | - | - | - | - | E | - | A | - | - | - | - | - | - | - | - | - | - | B | - |
| 11 |  |  |  | G | F | B | - | - | - | - | - | - | - | - | - | - | F | - | - | - | - | - | - | - | - | - | - | - | - | - | - |
| 12 |  | *Corapipo* |  | G | F | C | - | - | - | - | - | - | - | - | - | - | F | - | - | - | - | - | B | - | - | - | - | - | - | - | - |
| 13 |  |  |  | G | F | C | - | - | - | - | - | - | - | - | - | - | F | - | - | - | - | - | B | - | - | - | - | - | - | - | - |
| 14 |  | Piprini |  | D | - | B | - | - | - | - | - | - | - | - | - | - | - | - | - | - | - | - | - | - | - | - | - | - | - | - | - |
| 15 |  | *Xenopipo* |  | D | - | B | - | - | - | - | - | - | - | - | - | - | - | - | - | - | - | - | - | - | - | - | - | - | - | - | - |
| 16 |  |  |  | D | - | B | - | - | - | - | - | - | - | - | - | - | - | - | - | - | - | - | - | - | - | - | - | - | - | - | - |
| 17 |  | *Chloropipo* |  | D | - | B | - | - | - | - | - | - | - | - | - | - | - | - | - | - | - | - | - | - | - | - | - | - | - | - | - |
| 18 |  |  |  | D | - | B | - | - | - | - | - | - | - | - | - | - | - | - | - | - | - | - | - | - | - | - | - | - | - | - | - |
| 19 |  |  |  | D | - | B | - | - | - | - | - | - | - | - | - | - | - | - | - | - | - | - | - | - | - | - | - | - | - | - | - |
| 20 |  |  |  | D | - | B | - | - | - | - | - | - | - | - | - | - | - | - | - | - | - | - | - | - | - | - | - | - | - | - | - |
| 21 |  | *Pipra* |  | G | F | B | - | - | - | - | - | - | - | - | C | - | - | - | - | - | - | - | - | - | - | - | - | - | - | - | - |
| 22 |  |  |  | D | - | B | - | - | - | - | - | - | - | - | - | - | - | - | - | - | - | - | - | - | - | - | - | - | - | - | - |
| 23 |  |  |  | D | - | B | - | - | - | - | - | - | - | - | - | - | - | - | - | - | - | - | - | - | - | - | - | - | - | - | - |
| 24 |  | *Ceratopipra* |  | D | - | B | - | - | - | - | - | - | - | - | - | - | - | - | - | - | - | - | - | - | - | - | - | - | B | - | - |
| 25 |  | *Cryptopipo* |  | D | - | B | - | - | - | - | - | - | - | - | - | - | - | - | - | - | - | - | - | - | - | - | - | - | B | - | - |
| 26 |  | *Machaeropterus* |  | D | - | - | - | B | - | - | - | - | - | - | - | B | - | - | B | - | - | - | - | - | - | - | - | - | - | - | - |
| 27 |  |  |  | D | - | - | - | B | - | - | - | - | - | - | - | B | - | - | B | - | - | - | - | - | - | - | - | - | - | - | - |
| 28 |  |  |  | D | - | B | - | - | - | - | - | - | - | - | - | - | - | - | - | - | - | - | B | - | - | - | - | - | - | - | - |
| 29 |  | *Manacus* |  | D | - | B | - | - | - | - | - | - | - | - | - | - | - | - | - | - | - | B | B | - | - | B | - | - | - | - | - |

**Table S1 continued.**

|  |  |  |  |  |  |  | **Character** | | | | | | | | | | | | | | | | | | | | | | | | |
| --- | --- | --- | --- | --- | --- | --- | --- | --- | --- | --- | --- | --- | --- | --- | --- | --- | --- | --- | --- | --- | --- | --- | --- | --- | --- | --- | --- | --- | --- | --- | --- |
| **Node** |  | **Ancestor** |  | **1** | **2** | **3** | **4** | **5** | **6** | **7** | **8** | **9** | **10** | **11** | **12** | **13** | **14** | **15** | **16** | **17** | **18** | **19** | **20** | **21** | **22** | **23** | **24** | **25** | **26** | **27** | **28** |
| 30 |  |  |  | A | - | B | - | - | - | - | - | - | - | - | - | - | - | - | - | - | - | B | B | - | - | B | - | - | - | - | - |
| 31 |  |  |  | D | - | B | - | - | - | - | - | - | - | - | - | - | - | - | - | - | - | - | - | - | - | - | - | - | - | - | - |
| 32 |  | *Lepidothrix* |  | D | - | B | - | - | - | - | - | - | - | - | - | - | - | - | - | - | B | - | - | - | - | - | - | - | - | - | - |
| 33 |  |  |  | D | - | B | - | - | - | - | - | - | - | - | - | - | - | - | - | - | B | - | - | - | - | - | - | - | - | - | - |

**Table S2.** Example transition models for a hypothetical plumage patch character with four character states (ϕ = Absent, plus A, B, and D). The custom ERϕ model distinguishes asymmetric parameters for gaining or losing a plumage patch, versus all other transitions that indicate shifts in developmental schedule. Numbers indicate distinct parameters within each model.

| *Equal rates (ER)* | | | | |  |  | *ERϕ* | | | |
| --- | --- | --- | --- | --- | --- | --- | --- | --- | --- | --- |
|  | ϕ | A | B | D |  |  | ϕ | A | B | D |
| ϕ | - | 1 | 1 | 1 |  | ϕ | - | 1 | 1 | 1 |
| A | 1 | - | 1 | 1 |  | A | 2 | - | 3 | 3 |
| B | 1 | 1 | - | 1 |  | B | 2 | 3 | - | 3 |
| D | 1 | 1 | 1 | - |  | D | 2 | 3 | 3 | - |
| *Symmetric (SYM)* | | | | |  |  | *All rates different (ARD) (SYM)* | | | |
|  | ϕ | A | B | D |  |  | ϕ | A | B | D |
| ϕ | - | 1 | 2 | 3 |  | ϕ | - | 1 | 2 | 3 |
| A | 1 | - | 4 | 5 |  | A | 4 | - | 5 | 6 |
| B | 2 | 4 | - | 6 |  | B | 7 | 8 | - | 9 |
| D | 3 | 5 | 6 | - |  | D | 10 | 11 | 12 | - |

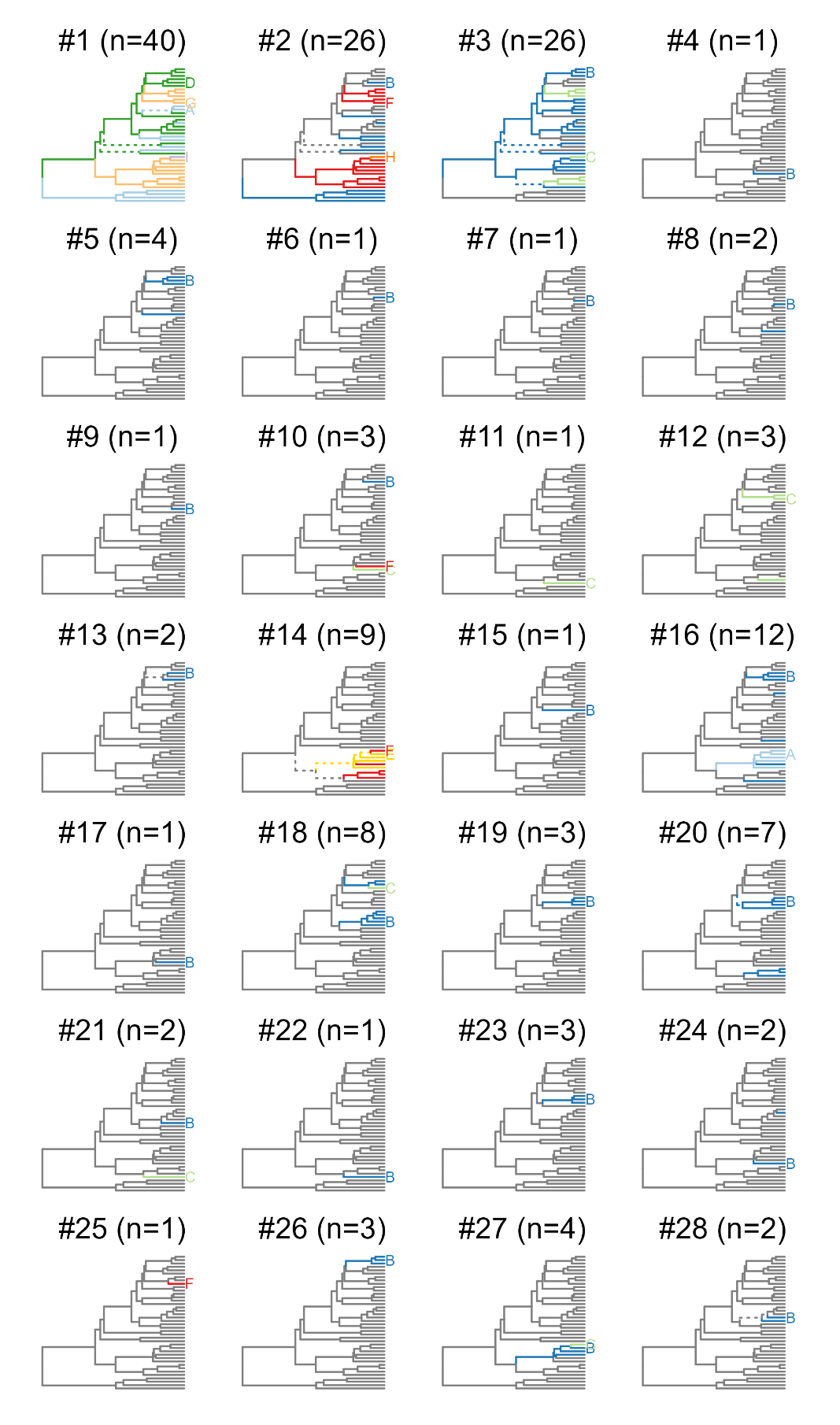

**Figure S2.** Plumage patch character histories. Panels are labeled by character number (with n = number of tip taxa where the character is present). Edges colored according to the maximum likelihood character states at each child node. Dashed lines indicate uncertainty (scaled likelihood <0.75).
